# Novel function acquired by the *Culex quinquefasciatus* mosquito D7 salivary protein enhances blood feeding on mammals

**DOI:** 10.1101/2020.01.24.918581

**Authors:** Ines Martin-Martin, Andrew Paige, Paola Carolina Valenzuela Leon, Apostolos G. Gittis, Olivia Kern, Brian Bonilla, Andrezza Campos Chagas, Sundar Ganesan, David N. Garboczi, Eric Calvo

## Abstract

Adult female mosquitoes require a vertebrate blood meal to develop eggs and continue their life cycle. During blood feeding, mosquito saliva is injected at the bite site to facilitate blood meal acquisition through anti-hemostatic compounds that counteract blood clotting, platelet aggregation, vasoconstriction and host immune responses. D7 proteins are among the most abundant components of the salivary glands of several blood feeding insects. They are members of a family of proteins that have evolved through gene duplication events to encode D7 proteins of several lengths. Here, we examine the ligand binding specificity and physiological relevance of two D7 long proteins, CxD7L1 and CxD7L2, from *Culex quinquefasciatus* mosquitoes, the vector of medical and veterinary diseases such as filariasis, avian malaria, and West Nile virus infections. CxD7L1 and CxD7L2 were assayed by microcalorimetry for binding of potential host ligands involved in hemostasis, including bioactive lipids, biogenic amines, and nucleotides/nucleosides. CxD7L2 binds serotonin, histamine, and epinephrine with high affinity as well as the thromboxane A2 analog U-46619 and several cysteinyl leukotrienes, as previously described for other D7 proteins. CxD7L1 does not bind any of the ligands that are bound by CxD7L2. Unexpectedly, CxD7L1 exhibited high affinity for adenine nucleotides and nucleosides, a binding capacity not reported in any D7 family member. We solved the crystal structure of CxD7L1 in complex with bound ADP to 1.97 Å resolution. The binding pocket for ADP is located between the two domains of CxD7L1, whereas all known D7s bind ligands either within the N-terminal or the C-terminal domains. We demonstrated that these two CxD7 long proteins inhibit human platelet aggregation in *ex vivo* experiments. CxD7L1 and CxD7L2 help blood feeding in mosquitoes by scavenging host molecules that promote vasoconstriction, platelet aggregation, itch, and pain at the bite site. The novel ADP-binding function acquired by CxD7L1 evolved to enhance blood feeding in mammals where ADP plays a key role in platelet aggregation.

## **1.** Introduction

*Culex quinquefasciatus* (Diptera: Culicidae) commonly known as the southern house mosquito, is a vector of medical and veterinary importance of filaria parasites, including *Wuchereria bancrofti* and *Dirofilaria immitis*^1, 2^ and avian malaria parasites (*Plasmodium relictum)*^3^. They also can transmit several arboviruses including Rift Valley fever, West Nile, St. Louis or Western equine encephalitis viruses^4, 5^. Adult female mosquitoes need to acquire vertebrate blood for egg development. During blood feeding, mosquito saliva is injected at the bite site and facilitates blood meal acquisition through anti-hemostatic compounds that prevent blood clotting, platelet aggregation and vasoconstriction as well as host immune responses^6^.

D7 proteins are among the most abundant components in the salivary glands of several blood feeding arthropods and are distantly related to the arthropod odorant-binding protein superfamily^7–10^. As mosquitoes adapted to consume different blood meals, D7 proteins evolved different biological activities to counteract the hemostatic response of their new vertebrate hosts^6^. The D7s belong to a multi-gene family that evolved through gene duplication events, resulting in long forms and truncated versions of a duplicated long form, known as short forms^8^. In addition to gene duplication, D7 proteins have undergone functional divergence, resulting in binding specialization with different affinities for host biogenic amines, as seen in *Anopheles gambiae* D7 short forms^10^. The D7 proteins act as kratagonists, binding and trapping agonists of hemostasis, including biogenic amines and leukotrienes (LT)^8, 11, 12^. The D7 long protein from *Anopheles stephensi* and intermediate D7 forms from the sand fly *Phlebotomus papatasi* have lost the capacity to bind biogenic amines but have evolved the capability to scavenge thromboxane A2 (TXA_2_) and LT^13, 14^, mediators of platelet aggregation and inflammation. Interestingly, an *Aedes aegypti* D7 long protein has a multifunctional mechanism of ligand binding: The N-terminal domain binds cysteinyl LT while the C-terminal domain shows high affinity to biogenic amines such as norepinephrine, serotonin, or histamine^10, 11^.

Many authors have studied this group of proteins since the first description of a D7 salivary protein in a blood feeding arthropod^15^. D7 proteins play a role in blood feeding function, mosquito physiology, and alter pathogen infection or dissemination^16–19^. Although the function of several mosquito D7 proteins including *An. gambiae* D7 short forms as well as the *Ae. aegypti* and *An. stephensi* long forms have been deciphered^10, 11, 13^, the role of *C. quinquefasciatus* D7 proteins remains unknown.

In this work, we expressed, purified, and biochemically characterized the two D7 long forms, L1 and L2, from *C. quinquefasciatus* salivary glands. We show the different affinities for biogenic amines and eicosanoids to CxD7L2 and discovered a new function for CxD7L1. CxD7L1 has a high affinity for adenosine 5′-monophosphate (AMP), adenosine 5′-diphosphate (ADP), adenosine 5′-triphosphate (ATP), and adenosine, which are essential agonists of platelet aggregation and act as inflammatory mediators that can prevent a successful bloodmeal. CxD7L1 showed no binding to biogenic amines or eicosanoids, that are previously described ligands for other D7 proteins^10, 11, 13^. We determined the crystal structure of CxD7L1 in complex with ADP and observed that the ADP binding pocket is located between the N-terminal and C-terminal domains. CxD7L1 is the first D7 protein to be shown to bind its ligands between the domains. We also show that CxD7L1 and CxD7L2 act as platelet aggregation inhibitors *ex vivo* supporting the hypothesis that the binding of ADP by CxD7L1 helped *C. quinquefasciatus* to evolve from blood feeding on birds, where serotonin plays a key role in aggregation, to blood feeding on mammals where ADP is a key mediator of platelet aggregation.

## 2. Results

### 2.1 Characterization of *Culex quinquefasciatus* CxD7L1 and CxD7L2

In previous studies^7, 8^, *Culex quinquefasciatus* salivary gland cDNA libraries were sequenced resulting in the identification of 14 cDNA clusters with high sequence similarity to the previously known two D7 long forms (D7clu1: AF420269 and D7clu12: AF420270) and a D7 short form (D7Clu32, AF420271). We compared the amino acid sequence of *C. quinquefasciatus* D7 long proteins with other well characterized mosquito and sand fly D7 members, whose function and structure have been solved. Exonic regions were conserved for all previously studied mosquito proteins (*Culex*, *Aedes* and *Anopheles*) where the first exon corresponds to a secretion signal peptide and the mature proteins are encoded by exons 2, 3, 4, and 5 (Fig. 1).

**Fig. 1.**
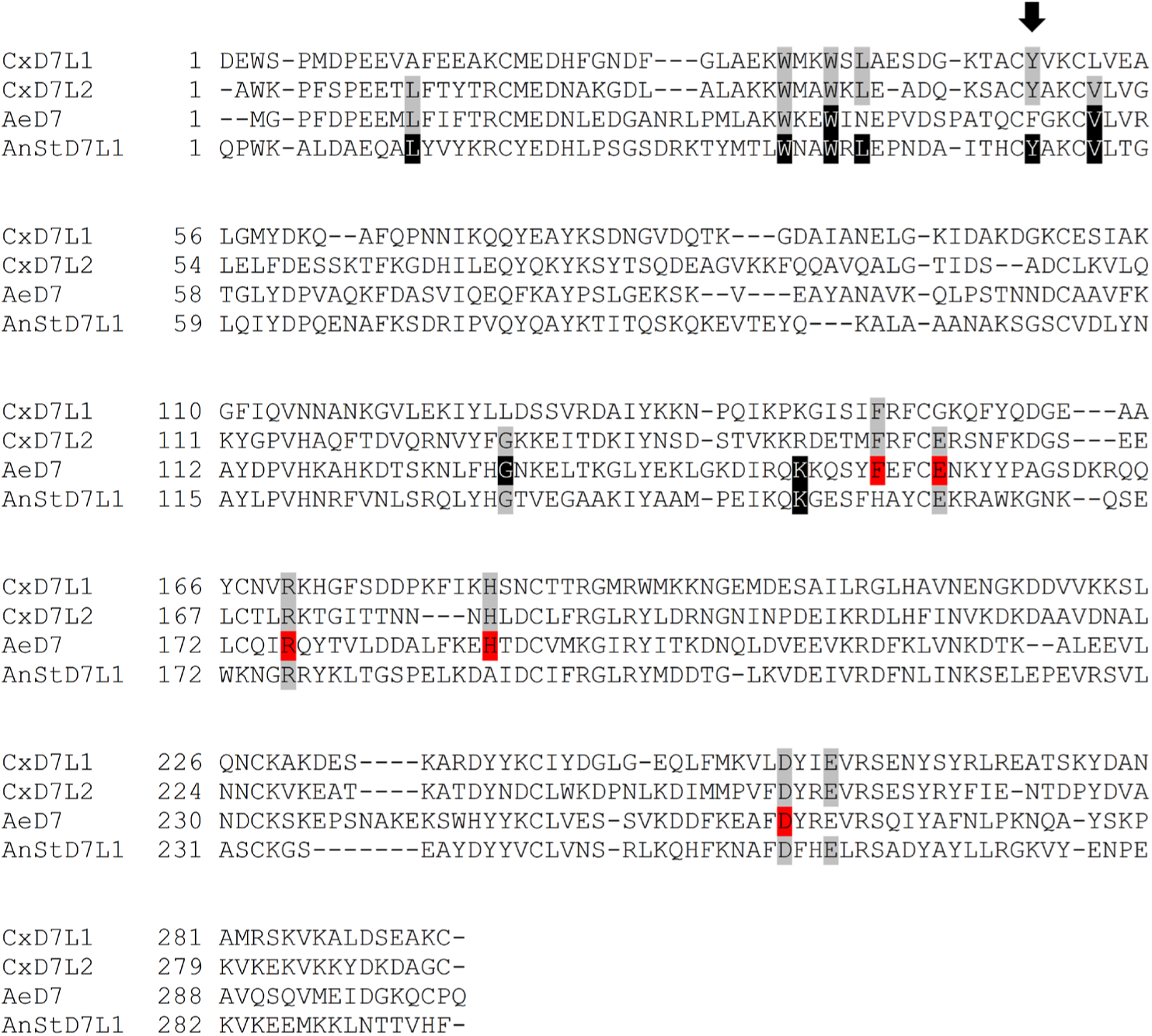
Multiple sequence alignment of *C. quinquefasciatus* D7 proteins and other related sequences. Comparison of *Culex* D7 long proteins: CxD7L1 (AAL16046) and CxD7L2 (AAL16047) with *Ae. aegypti* D7: AeD7 (PDB ID: 3DZT) and *An. stephensi* D7L1: AnStD7L1 (PDB ID: 3NHT). Sequences without a signal peptide were aligned with Clustal Omega and refined using BoxShade server. Black background shading represents amino acids involved in the eicosanoid binding of AeD7 and AnStD7L1^11, 13^. Red shading highlights amino acids involved in biogenic amine binding for AeD7^11^. Position K52, highlighted with an arrow, is involved in TXA_2_ binding^13^. Gray shading shows conserved residues of the amino acids involved in ligands binding.

We named *Culex quinquefasciatus* salivary long D7 proteins CxD7L1 (AAL16046) and CxD7L2 (AAL16047) and characterized them by gene expression analysis and immunolocalization. To determine the stage, sex, and tissue specificity of the D7 protein transcripts, qPCR experiments were performed on all four larval instars, pupae, whole male, whole female, female head and thorax, and female abdomen. We confirmed that both transcripts are only found in female adult stages with similar levels of expression and specifically located in the head and thorax of the mosquito, where the salivary glands are located. No amplification of *CxD7L1* and *CxD7L2* transcripts was found in the abdomen (Fig. 2a). These results confirmed that CxD7L1 and CxD7L2 expression is unique to the female salivary glands of *C. quinquefasciatus,* as previously shown in *Culex* and *Anopheles* mosquitoes^20, 21^.

**Fig. 2.**
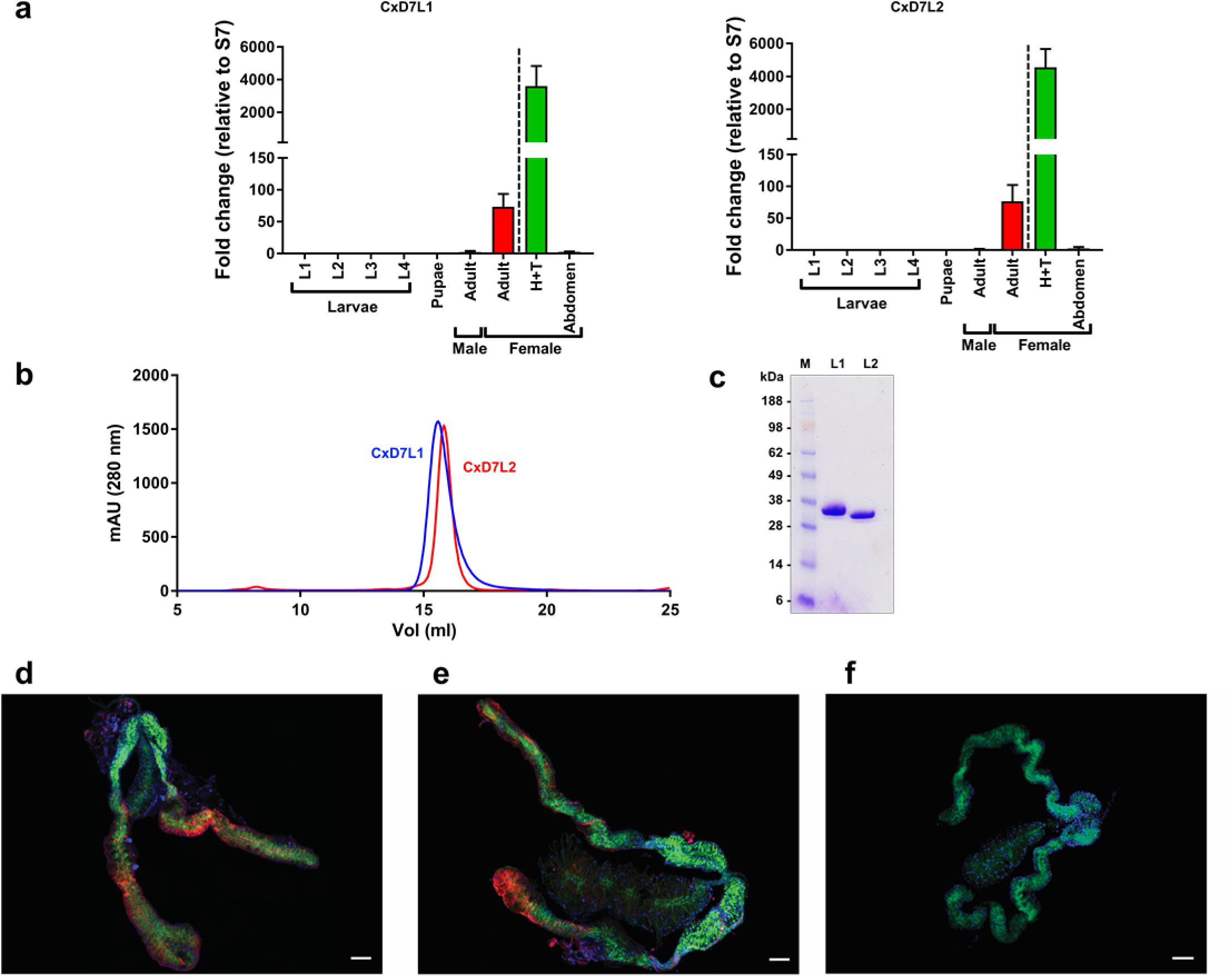
Characterization of *Culex quinquefasciatus* salivary long D7 proteins. (a) Gene expression analysis of *CxD7L1* and *CxD7L2* transcripts in different stages of *C. quinquefasciatus* mosquitoes. Relative abundance was expressed as the fold change using the 40S ribosomal protein S7 as the housekeeping gene. Larvae stage 1 (L1), larvae stage 2 (L2), larvae stage 3 (L3), larvae stage 4 (L4), pupae, male adult (reference sample), female adult, heads and thoraxes (H+T) and abdomens from female adult mosquitoes were analyzed separately. **(b)** Purification of CxD7L1 (blue line) and CxD7L2 (red line) by size exclusion chromatography using Superdex 200 Increase 10/300 GL column. **(c)** Coomassie-stained NuPAGE Novex 4-12% Bis-Tris gel electrophoresis of recombinant proteins CxD7L1 and CxD7L2 (1.5 µg). SeeBlue Plus2 Pre-stained was used as the protein standard (M). **(d** and **e)** Immunolocalization of CxD7L1 and CxD7L2 proteins in the salivary glands of *C. quinquefasciatus*. Salivary glands were incubated with rabbit IgG anti-CxD7L1 **(d)**, anti-CxD7L2 **(e)** and further stained with anti-rabbit IgG Alexa Fluor 594 antibody showed in red. Proteins of interest were localized in the medial and distal regions of the lateral lobes of *C. quinquefasciatus* salivary glands. As a control, salivary glands were incubated with anti-rabbit IgG AF594 alone **(f)**. Nucleic acids were stained by DAPI (blue) and the actin structure of salivary glands was stained using Phalloidin Alexa 488 (green). Scale bar = 50 µm.

To investigate the biochemical and biological activities of these proteins, CxD7L1 and CxD7L2 mature cDNA sequences were codon optimized for a eukaryotic cell expression system and engineered to contain a 6x-histidine tag in the C-terminal end followed by a stop codon. Both genes were subcloned into a VR2001-TOPO DNA cloning plasmid (Vical Inc) as described in Chagas *et al.*^22^. Recombinant CxD7L1 and CxD7L2 proteins were expressed in human embryonic kidney (HEK293) cells and purified by affinity and size exclusion chromatography (Fig. 2b). The identities of purified recombinant proteins were confirmed by N-terminal and liquid chromatography tandem mass spectrometry (LC/MS/MS sequencing). Both purified recombinant proteins migrated as single bands on Coomassie-stained precast polyacrylamide gels, and their apparent molecular weight (MW) in the gel corresponds to predicted MWs: 34.4 kDa and 34.8 kDa for CxD7L1 and CxD7L2, respectively (Fig. 2c). Immunogenicity of both proteins in their recombinant forms was maintained, as they were recognized by the purified IgG antibodies from a rabbit immunized against *C. quinquefasciatus* salivary gland extract (Supplementary Fig. 1a).

To perform immunolocalization experiments, specific antibodies against CxD7L1 and CxD7L2 were raised in rabbits. Because of the sequence similarity between these two proteins (34% identity), their antibodies showed cross-reactivity (Supplementary Fig. 1). To eliminate antibody cross-reactions and accurately identify D7 long form expression within salivary gland tissues, anti-CxD7L1 IgG was pre-adsorbed with CxD7L2 and anti-CxD7L2 IgG was pre-adsorbed with CxD7L1 (Supplementary Fig. 1). Using preabsorbed antibodies allowed us to accurately localize the *Culex* D7 long proteins within the female salivary glands. As shown in Figure 2d-f, CxD7L1 and CxD7L2 proteins are localized in the distal lateral and medial lobes of *C. quinquefasciatus* salivary glands, a pattern consistent with transcribed RNA of D7 long proteins in *Ae. aegypti* and *An. gambiae*^23, 24^.

### 2.2 *Culex quinquefasciatus* CxD7L1 binds adenine-nucleosides and nucleotides

Previous work demonstrated that members of the D7-related protein family can bind to biogenic amines and eicosanoids^10, 11, 13, 14^. Scavenging these proinflammatory and hemostatic mediators may have conferred an evolutionary adaptation to blood-feeding in mosquitoes. While *Culex* D7 proteins were first described in 2003^21^ and their transcripts were sequenced a year later^7^, their biological activity remains unknown. The binding abilities of CxD7L1 were tested with a wide panel of pro-hemostatic compounds including biogenic amines, nucleic acids, and proinflammatory lipids using isothermal titration calorimetry (ITC). In contrast to its D7 orthologs in *Aedes* and *Anopheles* mosquitoes, CxD7L1 does not bind biogenic amines such as serotonin, nor the pro-inflammatory lipids LTB_4_ and LTD_4_ or the stable analog of TXA_2_, U-46619 (Supplementary Fig. 2). However, CxD7L1 has evolved to bind adenine-nucleosides and nucleotides with high affinity (Table 1, Fig. 3), a novel function in a D7-related protein.

**Fig. 3.**
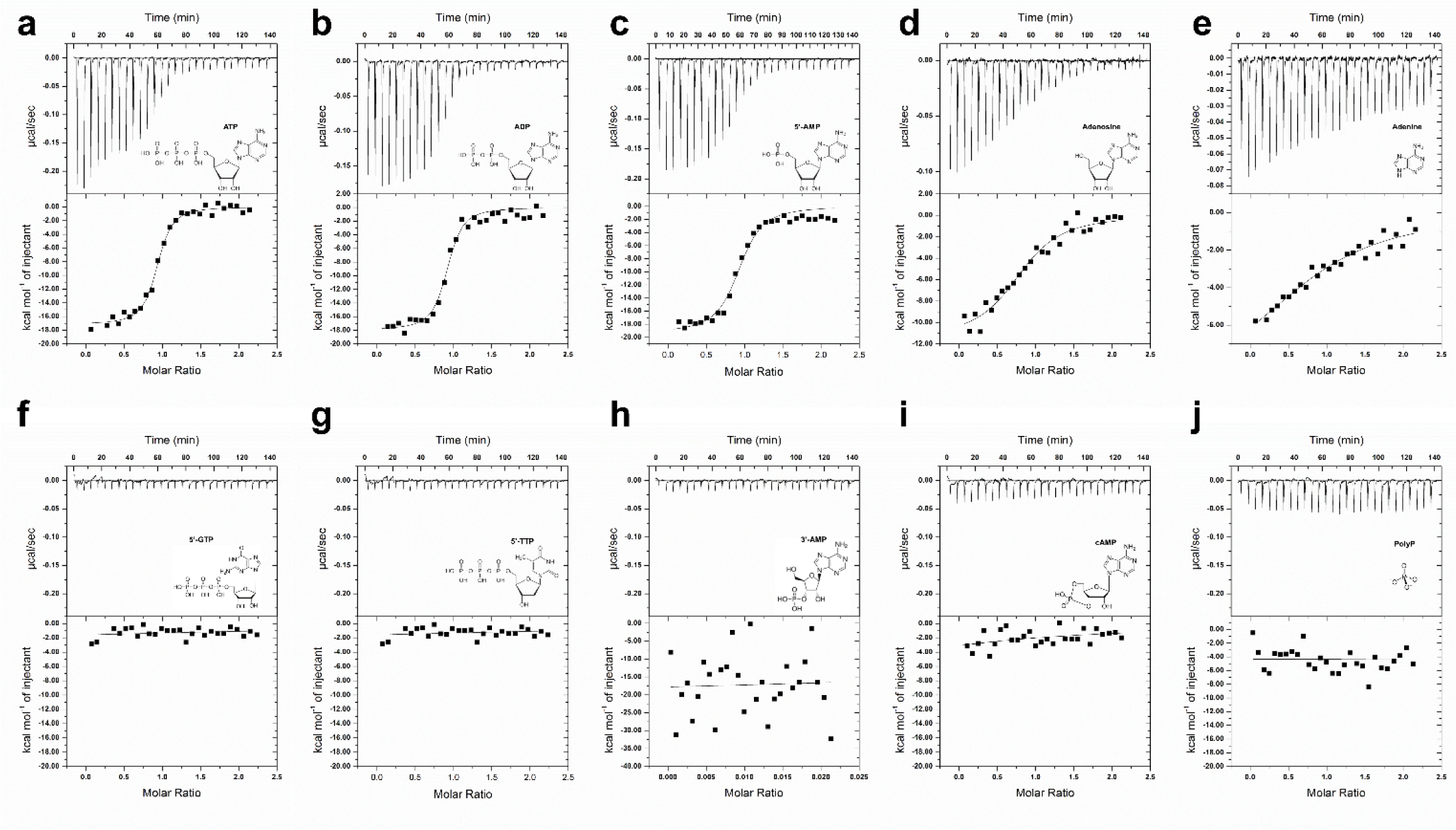
Binding of nucleosides and related molecules to CxD7L1 by isothermal titration calorimetry. Binding experiments were performed on a VP-ITC microcalorimeter. Assays were performed at 30 °C. The upper curve in each panel shows the measured heat for each injection, while the lower graph shows the enthalpies for each injection and the fit to a single-site binding model for calculation of thermodynamic parameters. Titration curves are representative of at least two measurements. Panels a-e show adenine nucleosides or nucleotides that bind CxD7L1: adenosine 5-triphosphate **(a)**, adenosine 5-diphosphate **(b)**, adenosine 5-monophosphate **(c)**, adenosine **(d)** and adenine **(e)**. In panels j-f other purine and pyrimidine nucleotides and related substances showed no binding to CxD7L1: guanosine 5-triphosphate **(f)**, thymidine 5-triphosphate **(g)**, adenosine 3-monophosphate **(h)**, cyclic adenosine monophosphate **(i)** and polyphosphate **(j)**. The insets show the names and chemical formulas for these compounds.

**Table 1.**
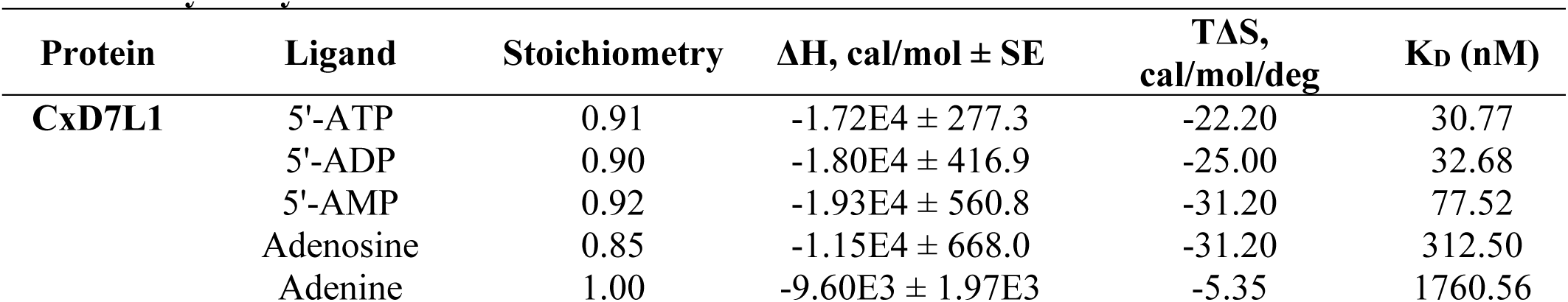
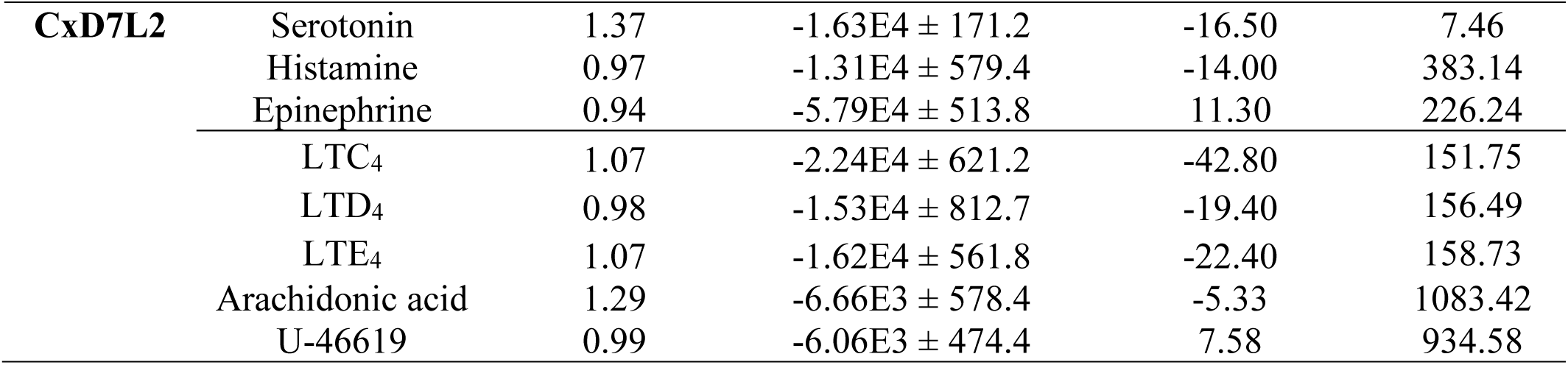
Thermodynamic parameters of Culex quinquefasciatus D7 proteins by isothermal.

| Protein | Ligand | Stoichiometry | $\Delta H$, cal/mol ± SE | $T\Delta S$, cal/mol/deg | $K_D$ (nM) |
| --- | --- | --- | --- | --- | --- |
| CxD7L1 | 5'-ATP | 0.91 | -1.72E4 ± 277.3 | -22.20 | 30.77 |
| | 5'-ADP | 0.90 | -1.80E4 ± 416.9 | -25.00 | 32.68 |
| | 5'-AMP | 0.92 | -1.93E4 ± 560.8 | -31.20 | 77.52 |
| | Adenosine | 0.85 | -1.15E4 ± 668.0 | -31.20 | 312.50 |
| | Adenine | 1.00 | -9.60E3 ± 1.97E3 | -5.35 | 1760.56 |
| **CxD7L2** | Serotonin | 1.37 | $-1.63\text{E}4 \pm 171.2$ | -16.50 | 7.46 |
| | Histamine | 0.97 | $-1.31\text{E}4 \pm 579.4$ | -14.00 | 383.14 |
| | Epinephrine | 0.94 | $-5.79\text{E}4 \pm 513.8$ | 11.30 | 226.24 |
| | LTC<sub>4</sub> | 1.07 | $-2.24\text{E}4 \pm 621.2$ | -42.80 | 151.75 |
| | LTD<sub>4</sub> | 0.98 | $-1.53\text{E}4 \pm 812.7$ | -19.40 | 156.49 |
| | LTE<sub>4</sub> | 1.07 | $-1.62\text{E}4 \pm 561.8$ | -22.40 | 158.73 |
| | Arachidonic acid | 1.29 | $-6.66\text{E}3 \pm 578.4$ | -5.33 | 1083.42 |
| | U-46619 | 0.99 | $-6.06\text{E}3 \pm 474.4$ | 7.58 | 934.58 |

Our biochemical characterization shows that CxD7L1 specifically binds the purine nitrogenous base adenine, its nucleoside (adenosine), and nucleotide derivates: AMP, ADP, and ATP, with the highest affinity to ATP and ADP (Fig. 3a-e). The binding is adenine-specific, as no binding was observed with other purine or pyrimidine nucleotides such as GTP or TTP (Fig. 3f-g). Although adenine is essential for binding, CxD7L1 did not bind to adenosine 3′-monophosphate (3’-AMP) or cyclic AMP (Fig. 3h-i), highlighting the importance of the phosphate group position in binding stabilization. Interaction between CxD7L1 protein and phosphate alone was ruled out as polyphosphate (sodium phosphate glass type 45) did not bind to the protein in ITC experiments (Fig. 3j). Furthermore, CxD7L1 did not bind to inosine (Supplementary Fig. 2), an intermediate metabolite in the purine metabolic pathway.

### 2.3 *Culex quinquefasciatus* CxD7L2 binds to serotonin, histamine, epinephrine, and eicosanoids

A detailed analysis of binding activities using ITC shows that CxD7L2 has comparable ligand binding capabilities as previously described in *Aedes* long and *Anopheles* long and short D7 proteins (Table 1, Fig. 4)^10, 11, 13^. CxD7L2 tightly binds serotonin (K_D_ = 7.5 nM) and other biogenic amines, including histamine and epinephrine, with lower affinities. It does not, however, bind norepinephrine. CxD7L2 also binds the cysteinyl leukotrienes, LTC_4_, LTD_4_, and LTE_4_ with a stoichiometry of 1:1 all with similar binding affinities (KD = 151.8 nM, 156.5 nM and 158.7 nM, respectively, Table 1, Fig. 4). CxD7L2 also binds arachidonic acid and U-46619, the stable analog of thromboxane A_2_, with lower affinities (K_D_ = 1083.42 nM and K_D_ = 934.6 nM, respectively) when compared to the cysteinyl LT. No binding to LTB_4_ was detected.

**Fig. 4.**
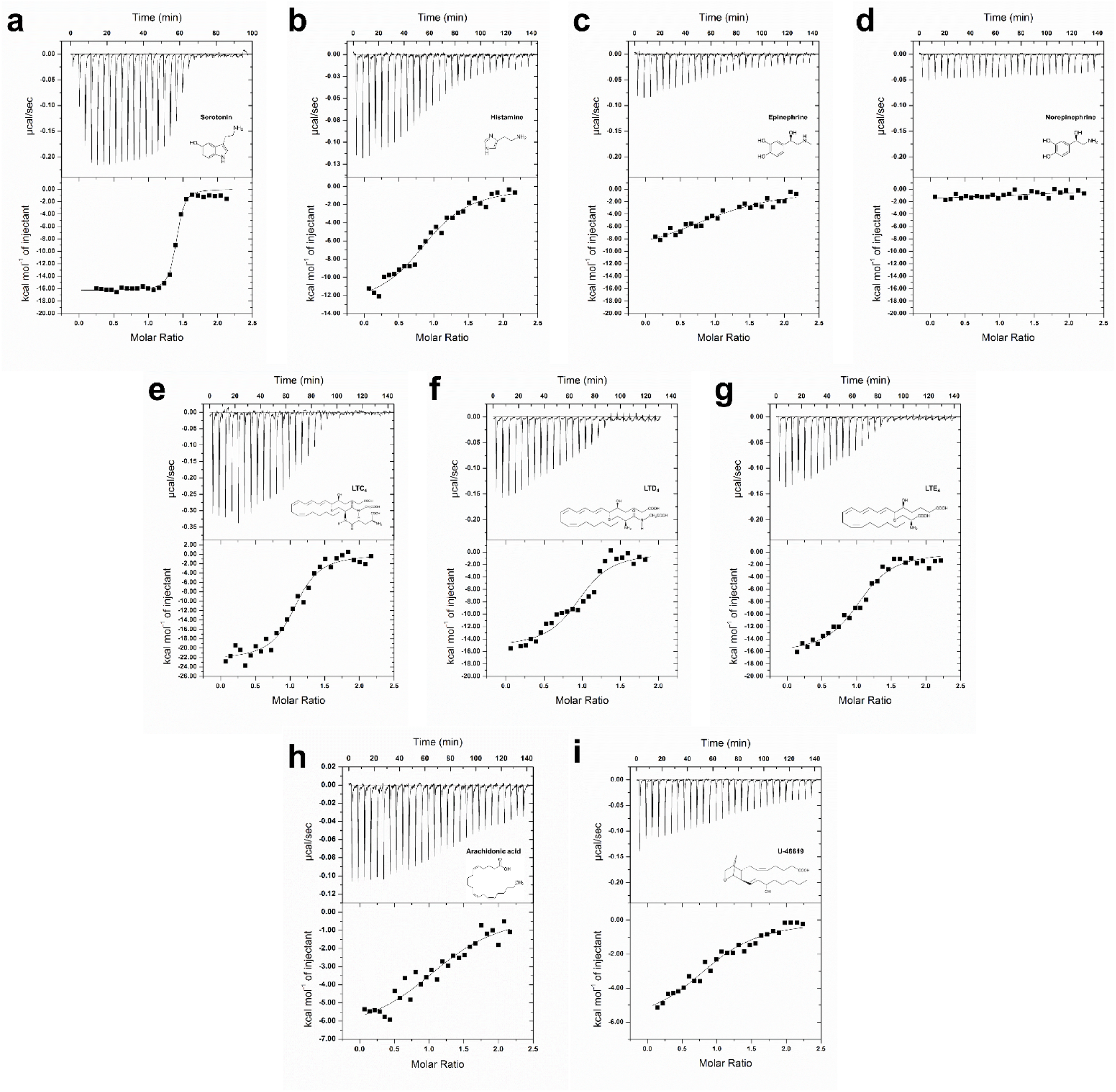
Binding of biogenic amines and eicosanoids to CxD7L2 by isothermal titration calorimetry. Binding experiments were performed on a VP-ITC microcalorimeter. The upper curve in each panel shows the measured heat for each injection, while the lower graph shows the enthalpies for each injection and the fit to a single-site binding model for calculation of thermodynamic parameters. Titration curves are representative of at least two measurements. Panels: serotonin **(a)**, histamine **(b)**, epinephrine **(c)** norepinephrine **(d),** LTC_4_ **(e),** LTD_4_ **(f),** LTE_4_ **(g)**, arachidonic acid **(h),** and TXA_2_ analog U-46619 **(i).** The insets show the names and chemical formulas for these compounds.

To gain insights into the mechanism of CxD7L2 binding to biogenic amines and eicosanoids, the N-terminal and C-terminal domains were independently cloned and expressed in *E. coli*. Only the C-terminal domain of CxD7L2 (CxD7L1-CT) was successfully purified and analyzed in parallel with the full-length protein by ITC. Similar to the full-length CxD7L2 protein, CxD7L2-CT binds to serotonin with high affinity (K_D_ = 1.5 nM, N = 1.06, ΔH = 4.31E4 ± 460 cal/mol; for CxD7L2-serotonin see Table 1). We concluded that CxD7L2-CT is responsible for the serotonin binding capacity displayed by the full-length protein. Since we were unable to produce the CxD7L2 N-terminal domain as a non-aggregated protein, a saturation study was designed to indirectly investigate the binding specificity of this domain. For this experiment, CxD7L2 protein was saturated with 50 µM serotonin (30 min pre-incubation) and titrated with LTD_4_ (in 50 µM of serotonin). The calculated binding parameters for CxD7L2 titrated with LTD_4_ in the absence or presence of serotonin remained similar (K_D_ = 156.8 nM, N = 0.93, ΔH = -2.21E4 ± 924.6 cal/mol; for CxD7L2-LTD_4_ see Table 1). These results demonstrate that lipids and biogenic amines bind to the CxD7L2 protein independently through different binding pockets, with lipids binding to the N-terminal pocket and biogenic amines to the C-terminal pocket, similar to the binding mechanism of AeD7 protein from *Ae. aegypti*^11^.

### 2.4 Crystal structure of *Culex quinquefasciatus* CxD7L1

To further characterize the mechanism of the novel adenine nucleoside/nucleotide D7 binding, we solved the crystal structure of CxD7L1 in complex with ADP. The structure of CxD7L1 was determined by molecular replacement using Phaser by employing separate, manually constructed search models for the N-terminal and C-domains based on the crystal structure of *Anopheles stephensi* AnStD7L1 (PDB ID: 3NHT). A crystal of CxD7L1 that belonged to I2_1_2_1_2_1_ space group and diffracted to 1.97 Å resolution was used to collect a data set (Table 2). The coordinates and structure factors have been deposited in the Protein Data Bank under the accession number 6V4C.

**Table 2.**
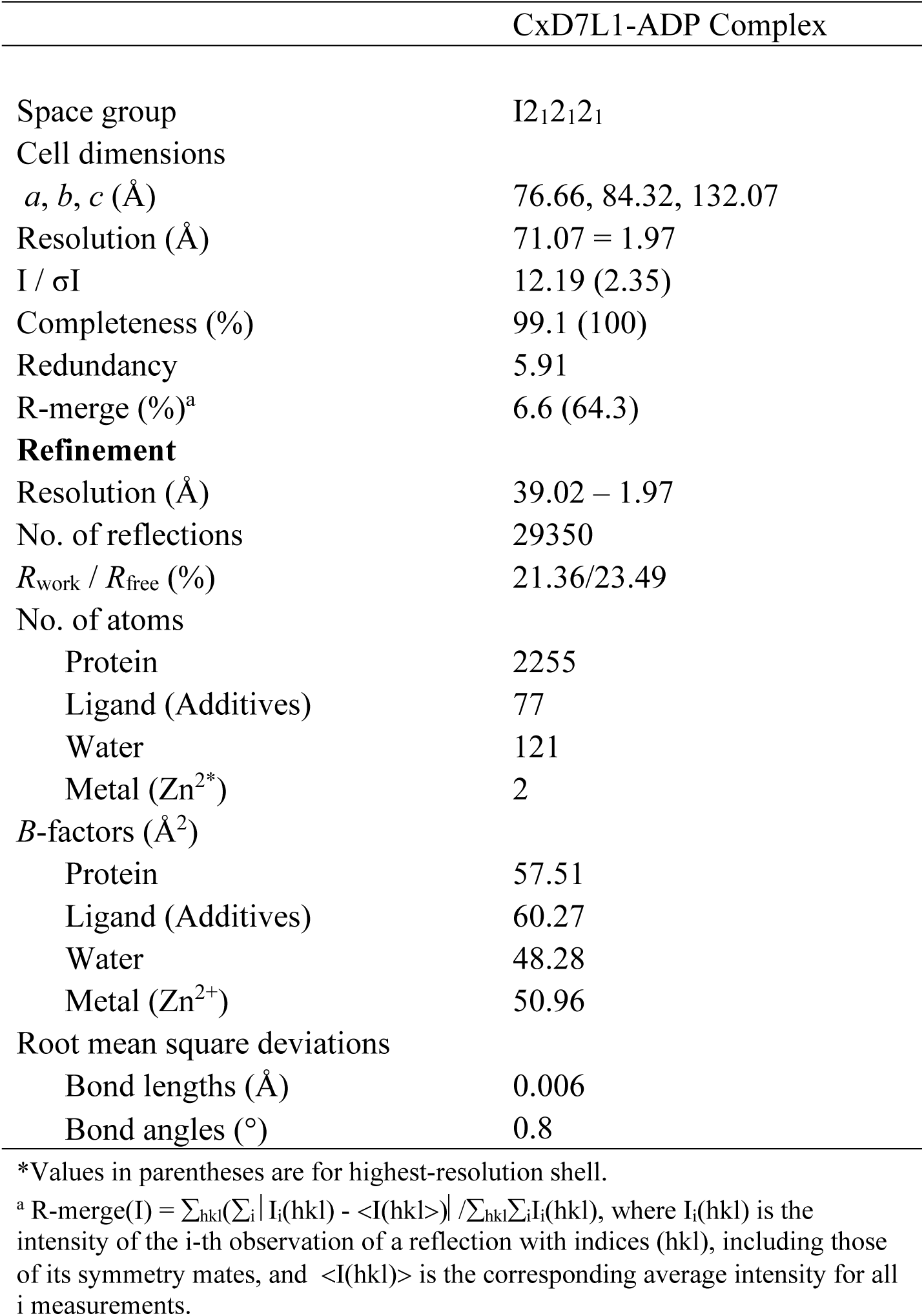
Data collection and refinement statistics.

The CxD7L1 protein fold consists of 17 helical segments stabilized by 5 disulfide bonds linking C18 with C51, C47 with C104, C154 with C186, C167 with C295 and C228 with C242 (Fig. 5a-b). The structure revealed that the ligand binding site is located between the N-terminal and C-terminal domains (Fig. 5a-e). All hydrogen bond donors and acceptors present in the adenine ring (N1, N3 and N7 are acceptors, and N6 is a donor) are interacting with the protein resulting in stable binding. The residues involved in binding ADP or stabilizing the binding pocket are R133, Y137, K144, K146, N265, Y266, S263, S267, and R271 (Fig. 5e). Residues Y137, K144 and Y266 bind to the adenine ring. The hydroxyl group of Y137 forms a bidentate hydrogen bond with the N6 and N7 of the adenine ring. The carbonyl oxygen of K144 forms a hydrogen bond with the amino nitrogen N6 of the adenine ring, while the NZ of K144 is involved in 2 hydrogen bonds, one with N1 from the adenine ring, and the other with the carbonyl oxygen of S263. It should be noted that the hydrogen bond with the carbonyl oxygen of S263 fixes NZ of the K144 in a position that allows it to bind the adenine ring. The amide nitrogen of Y266 binds N3 of the adenine ring and its side chain stacks partially on top of the base of ADP which provides a favorable van der Waals contribution to the CxD7L1-ADP interaction. As we go further along the ADP molecule, we find that S267 interacts strongly with and fixes the ribose ring of ADP with its hydroxyl group involved in 2 hydrogen bonds with both O2’ and O3’. In addition, the ribose oxygen O2’ forms a hydrogen bond with a water molecule and ND2 of N265 binds to O5’ of the sugar. Arginine 271 makes a hydrogen bond to N265 so that it is positioned favorably to engage in electrostatic interaction with the alpha phosphate. Lysine 146 is also in a location that can potentially be involved in electrostatic interaction with the alpha phosphate. Arginine 133 forms 2 salt bridges with the beta phosphate of ADP, with NH1 and NH2 of R133 binding to O1B and O3B of ADP respectively, which may explain the similar binding affinities between ATP and ADP and the lower affinity of AMP, which lacks the beta phosphate.

**Fig. 5.**
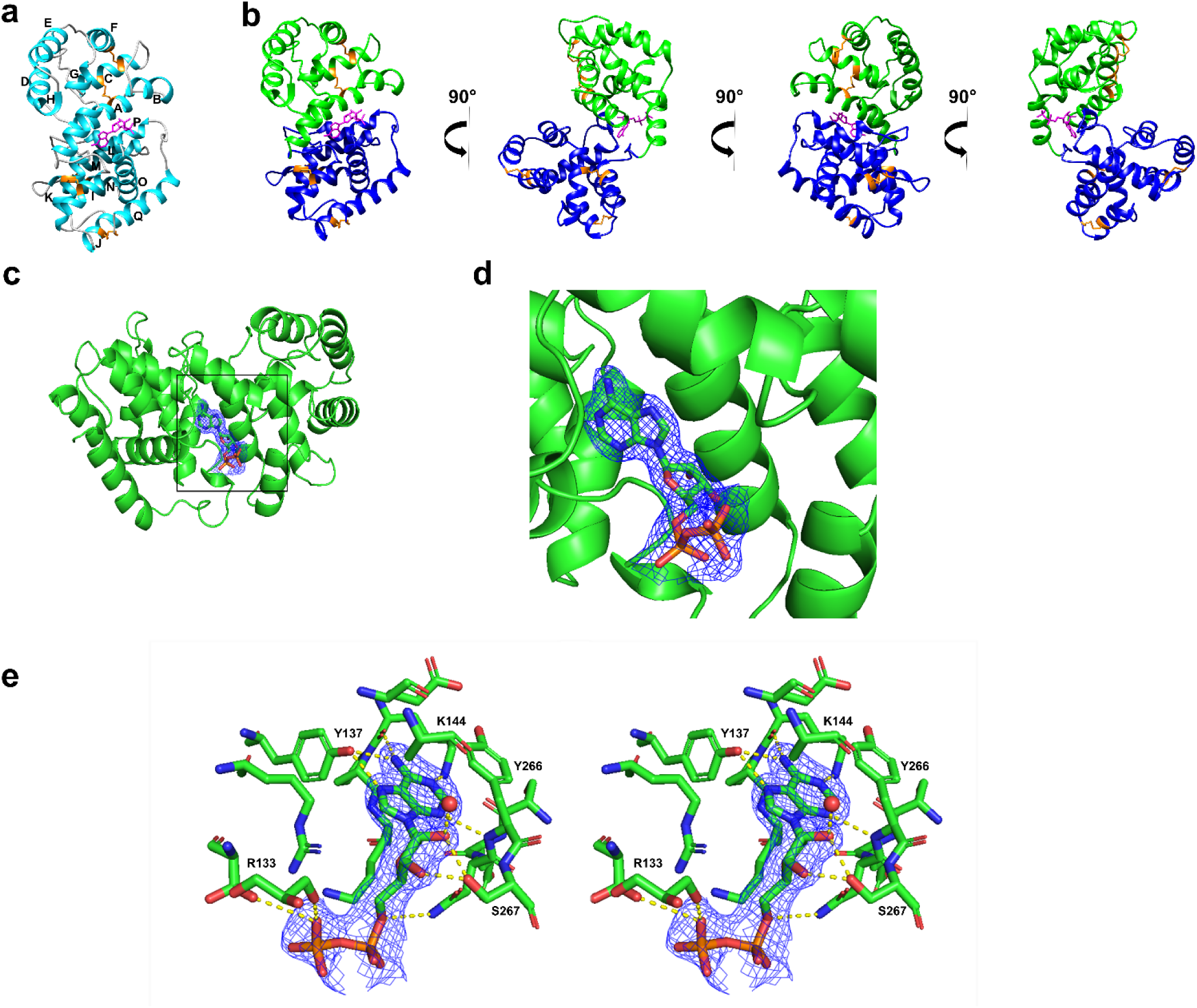
Structure of CxD7L1 in complex with ADP. (a) Ribbon representation of CxD7L1-ADP structure. The 17 α-helices are labelled A-Q. **(b)** Several views of CxD7L1 differing by rotations of 90 degrees around the y-axis. N-terminal and C-terminal are colored in blue and green, respectively. ADP is shown as a stick model in magenta and disulfide bonds in orange. **(c)** Electron density covering ADP. CxD7L1 protein is colored in green. Inset is shown in **(d)**. Amino acid residues of CxD7L1 involved in ADP binding are colored in green **(e)**. Stereo view of the binding pocket of the CxD7L1-ADP complex showing the 2*F_o_ – F_c_* electron density contoured at 1 σ covering the ligand. Hydrogen bonds are colored in yellow.

Although the superposition of structures of CxD7L1, AeD7 (PDB:3DZT), and AnStD7L1 (PDB:3NHT) showed a similar overall structure (Fig. 6a), the protein sequences only share 20% amino acid identity and some of the essential residues involved in the lipid and biogenic amine binding are missing in CxD7L1 (Fig. 1 and Supplementary Fig. S3). Moreover, CxD7L1 showed a completely different electrostatic surface potential (Coulombic Surface Coloring generated by Chimera software) when compared to *Ae. aegypti* D7L and *An. stephensi* AnStD7L1, which may contribute to the differences in their binding capacity. The amino acids that constitute the ADP binding pocket in CxD7L1 create a strongly negative surface, showing an inverted pattern of amino acid charges that completely change the nature of the binding pockets (Fig. 6b). The residues involved in ADP binding were not conserved in other D7 homologs (Fig. 1).

**Fig. 6.**
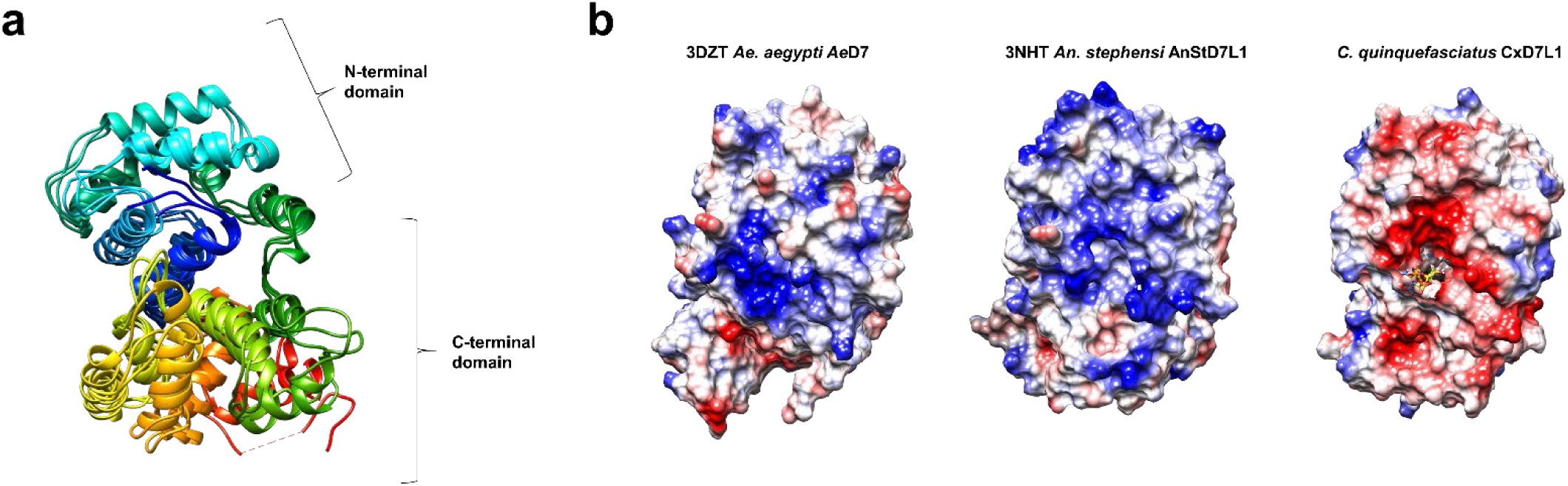
Multiple sequence superposition and electrostatic potential of *Culex* D7 proteins and other related sequences. (a) Superposition of CxD7L1, *Ae. aegypti* AeD7 (PDB ID: 3DZT) and *An. stephensi* AnStD7L1 (PDB ID: 3NHT) shows a similar overall helix structure. Rainbow coloring pattern shows the N-terminal in blue and the C-terminal in red. **(b)** Electrostatic potential of 3DZT, 3NHT and CxD7L1 generated by Coulombic Surface Coloring (Chimera software) with blue being positive and red being negative. ADP is represented as a stick model.

Although most of the residues were present in D7 long proteins from *Culex tarsalis* (Supplementary Fig. S3) no experimental data is available showing that D7L1 from this mosquito retains the ADP binding capacity.

### 2.5 *Culex quinquefasciatus* CxD7L1 and CxD7L2 play a role in platelet aggregation

Because CxD7 long forms bind platelet aggregation agonists such as ADP, serotonin, or the TXA_2_ analog U-46619, we examined their ability to interfere with platelet aggregation in *ex vivo* experiments. At low concentrations of collagen (1 µg/mL), we saw the classical collagen induction trace, where there is a delay of the platelet shape change due to the release of secondary mediators and observed as the initial decrease of light transmittance. There was a clear dose-dependent inhibition of platelet aggregation by both CxD7L1 and CxD7L2 (Fig. 7a). Neither CxD7L1 nor CxD7L2 interfered with platelet aggregation induced by high doses of either collagen (Fig. 7b) or convulxin (Fig. 7c), an agonist of the platelet GPVI collagen receptor which induces platelet aggregation independently of secondary mediators.

**Fig. 7.**
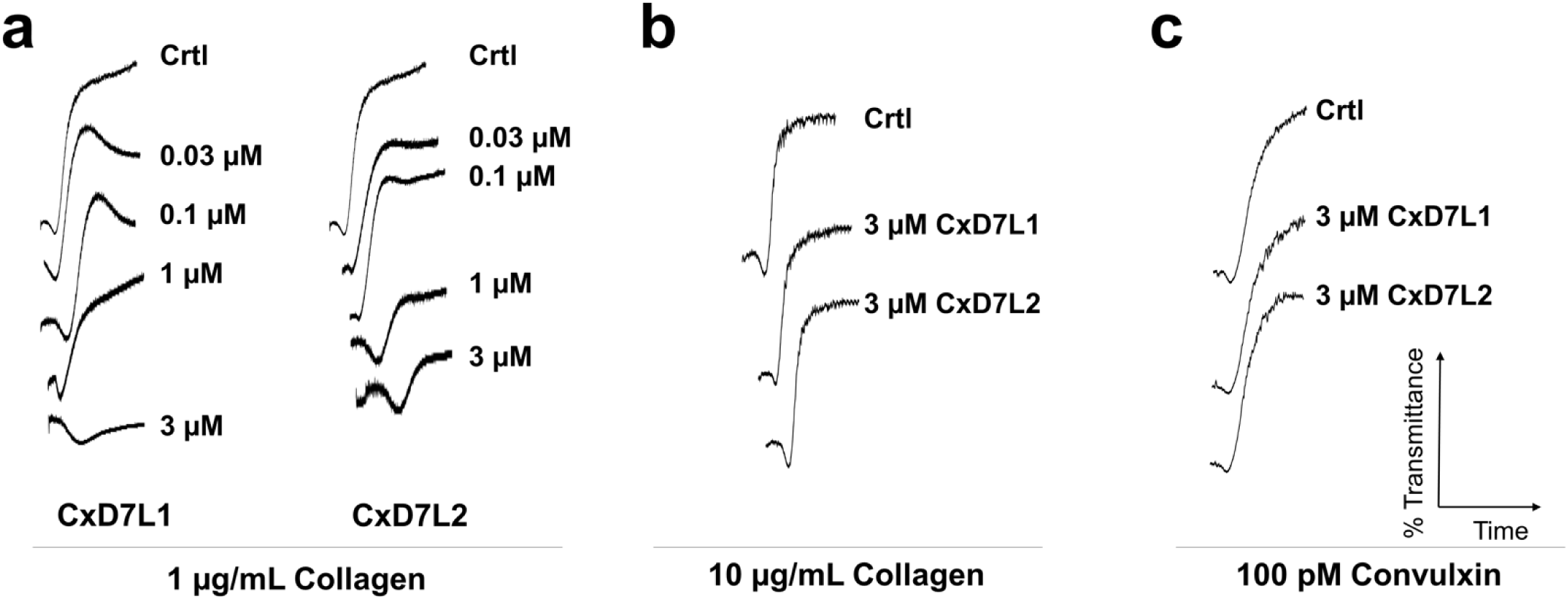
Effect of CxD7L1 and CxD7L2 on platelet aggregation induced by collagen or convulxin. Prior to the addition of the agonist, platelet-rich human plasma was incubated for 1 minute with either PBS (Crtl) or with the recombinant proteins at the concentrations shown. Aggregometer traces were measured at 37 °C from stirred platelets suspensions on a Chrono-Log platelet aggregometer model 700 for 6 min. An increase of light transmittance over time indicates platelet aggregation. **(a)** CxD7L1 and CxD7L2 concentration-dependent inhibition of platelet aggregation induced by low doses of collagen (1 μg/mL). CxD7L1 and CxD7L2 failed to inhibit platelet aggregation induced by **(b)** high doses of collagen (10 µg/mL) and **(c)** GPVI agonist convulxin (100 pM).

We also investigated the anti-platelet aggregation activity of CxD7L1 and CxD7L2 using ADP as an agonist. ADP plays a role in the initiation and extension of the aggregation cascade. In our studies, different concentrations of ADP were used as an agonist. When ADP was added at concentrations below the threshold for platelet aggregation (0.5 µM), only platelet shape change was observed (control trace, Fig. 8a). Preincubation of platelets with CxD7L1 prevented this shape change. With higher doses of ADP (1 µM), platelet aggregation was inhibited in the presence of 3 µM CxD7L1 (Fig. 8a). At high doses of ADP (10 µM), 3 µM of CxD7L1 was insufficient to inhibit platelet aggregation, confirming the nature of the inhibition by scavenging the mediator. The addition of CxD7L2 did not show any effect in aggregation initiated via ADP at any dose, confirming that CxD7L2 does not target ADP (Fig. 8a).

**Fig. 8.**
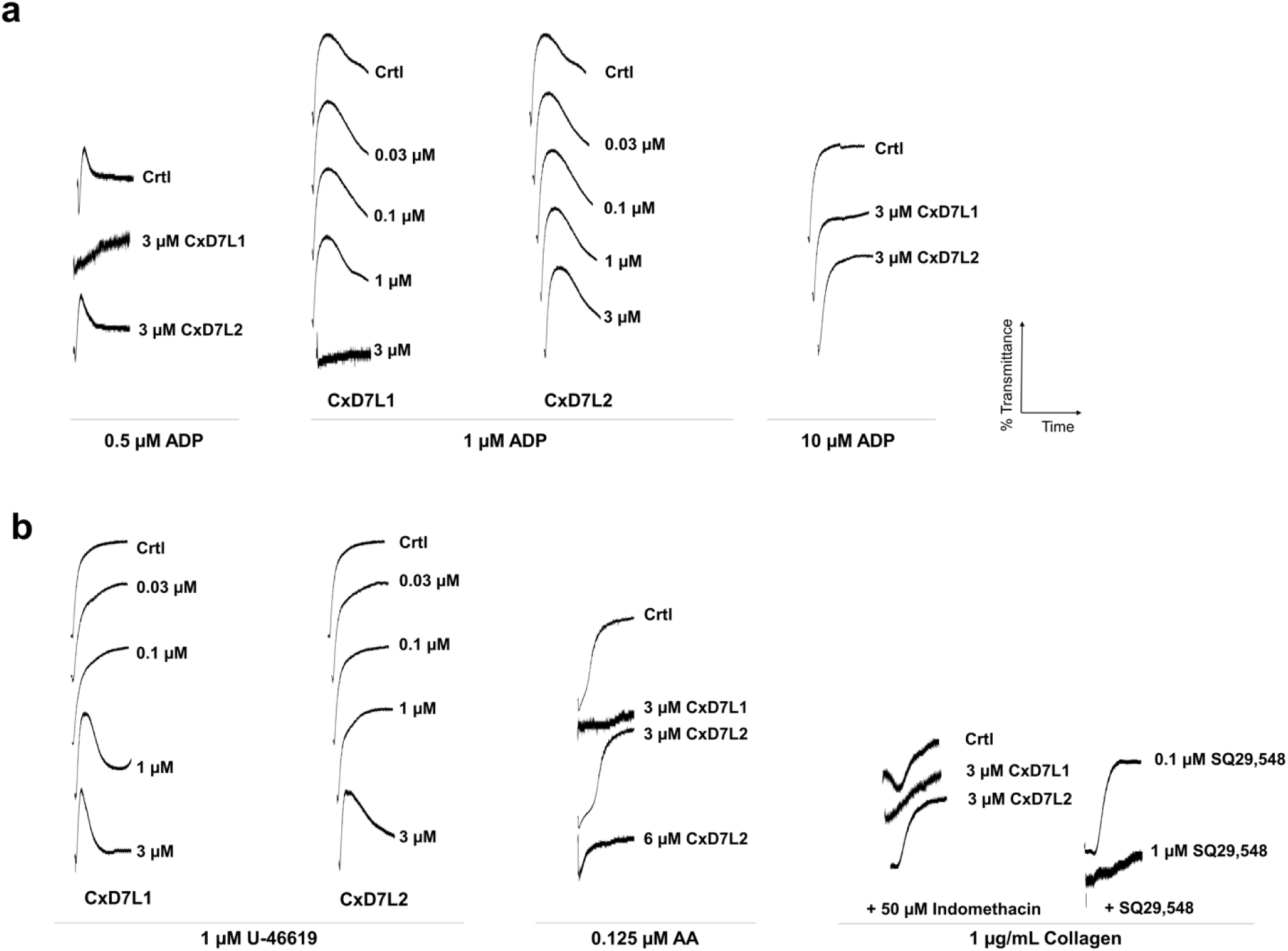
Effect of CxD7L1 and CxD7L2 on platelet aggregation induced by secondary mediators. Prior to the addition of the agonist, platelet-rich human plasma was incubated for 1 min either with PBS (Crtl) or with the recombinant proteins, or SQ29,548 at the concentrations shown. Aggregometer traces were measured at 37 °C from stirred platelets suspensions on a Chrono-Log platelet aggregometer model 700 for 6 min. An increase of light transmittance over time indicates platelet aggregation. **(a)** Platelet aggregation traces using different concentrations of ADP (0.5 µM, 1 µM and 10 µM) as aggregation agonist. **(b)** Platelet aggregation traces using 1 µM U-46619, 0.125 µM arachidonic acid (AA) or low collagen concentration (1 µg/mL).

We also used U-46619, the stable analog of TXA_2_ and widely accepted for platelet aggregation studies^13, 14, 26, 27^. When platelets are activated, TXA_2_ is synthesized from arachidonic acid released from platelet membrane phospholipids. TXA_2_ is an unstable compound and cannot be evaluated directly as a platelet aggregation agonist *ex vivo*. CxD7L2 inhibited U-46619-induced platelet aggregation in a dose-dependent manner. However, platelet shape change requires minimal concentrations of TXA_2_, and it was not prevented by CxD7L2 (Fig. 8b). Shape change was only abolished in the presence of 1 µM SQ29,548, a specific antagonist of the TXA_2_ receptor (Fig. 8b). This result is supported by our biochemical data showing that CxD7L2 binds directly to U-46619 *in vitro* (Fig. 4h). However, we do not know whether this binding is retained *in vivo*.

To verify that this protein binds the biological active TXA_2_ *ex vivo*, we induced platelet aggregation with its biosynthetic precursor, arachidonic acid, so that TXA_2_ would be released by platelets. CxD7L2 inhibited platelet aggregation induced by arachidonic acid only at high doses of protein (6 µM, Fig. 8b), most likely due to the low binding affinity observed for U-46619 and arachidonic acid (Table 1). To further investigate whether this effect was a result of a direct sequestering of TXA_2_ by CxD7L2, we pre-incubated platelets with indomethacin, a cyclooxygenase-1 inhibitor, that prevents TXA_2_ biosynthesis. We observed almost no inhibition of low dose collagen-induced platelet aggregation in the presence of CxD7L2 (Fig. 8b), indicating that the anti-platelet aggregation activity of CxD7L2 is mediated by TXA_2_ binding.

CxD7L1 inhibits platelet aggregation induced by U-46619 in a dose-dependent manner (Fig. 8b). CxD7L1 does not bind U-46619 as shown by microcalorimetry (Supplementary Fig. S2), but it tightly binds ADP (Fig. 3b, Table 1). Platelet aggregation triggered by U-46619, arachidonic acid, and low doses of collagen is highly dependent on ADP^28^. As a confirmation of this dependence, CxD7L1 inhibits platelet aggregation stimulated by either U-46619 or arachidonic acid as effectively as the antagonist of the TXA_2_ receptor SQ29,548. CxD7L1 also prevented aggregation initiated by low dose of collagen in indomethacin-treated platelets (Fig. 8b).

Serotonin acts as a potentiator of platelet agonists such as ADP or collagen. Alone, serotonin can initiate platelet aggregation, but in the absence of a more potent agonist, the platelets eventually disaggregate (Fig. 9a). CxD7L2 tightly binds serotonin (Fig. 4a). Therefore, the initiation of aggregation produced by serotonin was completely abolished in the presence of equimolar concentrations of the recombinant protein (Fig. 9a). However, when a higher dose of serotonin was used (10 μM), CxD7L2 was unable to sequester all the serotonin, resulting in no observed inhibition of platelet aggregation (Fig. 9a). When serotonin and low doses of collagen were used as aggregation agonists, CxD7L1 partially prevented aggregation, presumably due to its ADP binding, while CxD7L2-serotonin binding resulted in full inhibition of platelet aggregation (Fig. 9b). Serotonin also potentiated aggregation initiated by low doses of ADP (Fig. 9c). When platelets were incubated with CxD7L2, the synergistic effect of serotonin and ADP in platelet aggregation was abolished (Fig. 9c). CxD7L1, as a potent ADP-binder, completely abrogated platelet aggregation initiated by serotonin and ADP combined. In addition, CxD7L2 partially prevented aggregation initiated by serotonin and epinephrine (Fig. 9d).

**Fig. 9.**
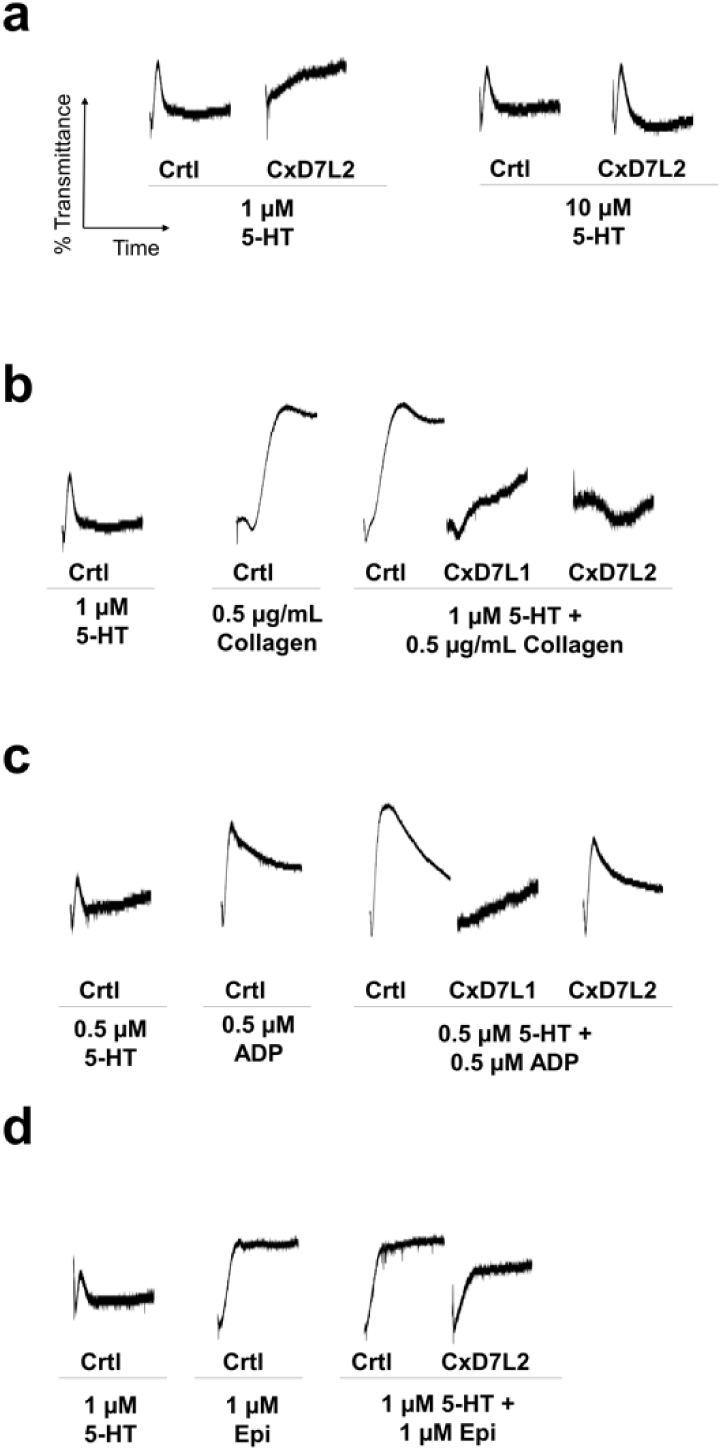
Effect of CxD7L1 and CxD7L2 on platelet aggregation induced by serotonin alone or in combination with collagen, ADP, or epinephrine. Prior to the addition of the agonist, platelet-rich human plasma was incubated for 1 minute either with PBS (Crtl) or with the recombinant proteins at the concentrations shown. Aggregometer traces were measured at 37 °C from stirred platelets suspensions on a Chrono-Log platelet aggregometer model 700 for 6 min. An increase of light transmittance over time indicates platelet aggregation. **(a)** Platelet aggregation traces using different concentrations of serotonin (5-HT) (1 µM and 10 µM) as aggregation agonist. **(b)** Platelet aggregation traces using 5-HT in combination with collagen, **(c)** ADP or **(d)** epinephrine (Epi).

## 3. Discussion

An arthropod blood feeding event can be considered as a battle between the need of the arthropod to acquire blood and the vertebrate host response to prevent blood loss. The outcome of this battle determines whether the arthropod can complete its life cycle, making a successful blood feeding event a crucial process for the fate of the invertebrate. During a bite, arthropod salivary proteins are injected into the host skin to counteract host hemostatic mediators. In this work, we characterized the structure and function of the salivary D7 long proteins from *C. quinquefasciatus* mosquitoes and described a novel mechanism of platelet aggregation inhibition for a D7 salivary protein.

CxD7L1 and CxD7L2 were found to be expressed in the distal-lateral and medial lobes of *C. quinquefasciatus* salivary glands. Salivary proteins have been shown to accumulate in the salivary glands forming distinct spatial patterns^23^. Although the relevance of distinct protein localization is not yet well understood, it supports the hypothesis of functionally-distinct regions within mosquito salivary glands. Salivary proteins related to sugar-feeding, nectar-related digestion, and bactericidal functions are localized in the proximal-lateral lobes, while proteins involved in blood-feeding, such as CxD7L1 and CxD7L2, are localized in the medial or distal-lateral lobes. More research is required to understand the implications of the salivary protein compartmentalization and viral infection of the glands.

D7 proteins are widely distributed in the saliva of hematophagous Nematocera, including mosquitoes, black flies, biting midges, and sand flies^8^. D7 salivary proteins antagonize the hemostasis mediators through a non-enzymatic, non-receptor-based mechanism by binding and sequestering several host hemostasis mediators^8, 10, 11, 13, 14^. This mechanism of action requires a high concentration of salivary protein at the bite site. As D7 proteins bind their ligands in a 1:1 stoichiometric ratio, they must be in equimolar concentrations with the mediators, which range from 1-10 µM for histamine, serotonin, or ADP^8^. This may explain why D7 salivary proteins are one of the most abundant components of the salivary glands.

Biogenic amines play important physiological roles in host hemostasis. Serotonin is released from platelet granules upon activation and acts as a weak platelet aggregation agonist. Serotonin and histamine increase vascular permeability and induce host sensations of pain and itch^29^. The catecholamines norepinephrine and epinephrine stimulate vasoconstriction by directly acting on adrenoreceptors^12^. Binding of biogenic amines by mosquito D7 proteins has been previously reported in the literature, highlighting the importance of removing these mediators at the bite site ^10, 11, 30^. Binding affinities for the different amines vary, as D7 proteins have become highly specialized for specific ligands^10, 11, 13, 14^. CxD7L2 tightly binds serotonin and epinephrine in the same range as the short D7 proteins from *An. gambiae* and AeD7 from *Ae. aegypti*^10, 11^. However, it showed lower affinity for histamine and did not bind norepinephrine. Like AeD7 from *Ae. aegypti*^11^, CxD7L2 is multifunctional and was able to bind biolipids through its N-terminal domain and biogenic amines through its C-terminal domain, as confirmed by ITC experiments. CxD7L2 binds cysteinyl leukotrienes (LTC_4_, LTD_4_, and LTE_4_) with similar affinities. Cysteinyl leukotrienes are potent blood vessel constrictors and increase vascular permeability^31^. The cysteinyl residue appears to play a role in lipid binding, as calorimetry experiments with lipids lacking a cysteinyl residue such as LTB_4_ showed no binding. Residues involved in bioactive lipid binding were conserved between CxD7L2 and the D7 proteins from *An. stephensi* and *Ae. aegypti* (AnStD7L1 and AeD7). Interestingly, a tyrosine residue at position 52 is present in *Culex* D7 long proteins and has been correlated to the ability to stabilize the binding of the TXA_2_ mimetic (U-46619) in *An. stephensi*^13^. This residue is absent in the *Ae. aegypti* D7 protein that does not bind U-46619^11^. This might explain the ability of CxD7L2 to bind cysteinyl leukotrienes and U-46619. Additionally, several residues known to be involved in the biogenic amine-binding were conserved in *Culex* D7 long proteins, for which the biogenic amine binding capability of CxD7L2 may be accounted.

Although CxD7L1 retains some amino acids involved in biogenic amine or lipid binding, ITC data showed that this protein lacks binding capacities typical of D7 proteins. Rather, CxD7L1 binds adenine nucleosides and nucleotides. Our crystallographic data clearly confirms our binding results. The nature of the binding pocket demonstrates specificity for the adenine ring. The hydrogen bonds between the adenine ring and residues Y137, K144, and Y266 determine the specificity for adenine and the lack of binding to other nucleotides with other nitrogenous bases (5’-GTP, 5’-TTP). Similarly, S267 and N265 of CxD7L1 are involved in binding to the ribose, which is possible when the phosphate group occupies position 5’ but not position 3’ or the cyclic form, as shown by calorimetry experiments. Arginine 133 binds to the oxygen of the beta phosphate of ADP which may explain the similar binding affinities for both ATP and ADP while affinity for AMP is lower as it lacks the beta phosphate.

CxD7 proteins scavenge biogenic amines, LTs, and ADP released at the bite site, and thus prevent hemostasis by inhibiting several simultaneous signaling cascades. Here, we have focused on their contributions in preventing platelet aggregation. Platelet aggregation occurs within seconds of tissue injury, restricting blood flow and creating a platelet plug that reduces blood feeding success. Exposure of circulating platelets to collagen from the subendothelial matrix or thrombin leads to the formation of a platelet monolayer that supports subsequent adhesion of activated platelets to each other^12, 32^. At low concentrations of collagen, ADP and TXA_2_ play an important role on the extension and amplification step of the platelet plug formation. Upon platelet activation, mediators secreted by platelets bind to G protein-coupled receptors in platelet membranes, rapidly amplifying the aggregation signal in a positive feedback response^33^.

However, at high concentrations, collagen acts as a strong agonist of the GPVI receptor on platelet surface, which induces platelet aggregation in an independent manner of ADP or TXA_2_ secretion^32^. Both CxD7L1 and CxD7L2 proteins showed a potent inhibitory effect on platelet aggregation, explained by distinct mechanisms. CxD7L2 inhibits platelet aggregation in the classical mechanism observed in other eicosanoid-scavenging salivary proteins^13, 14, 26, 34, 35^. CxD7L2 inhibits low dose collagen-induced platelet aggregation in a dose dependent manner but did not affect aggregation induced by high doses of collagen or convulxin. These findings indicate that CxD7L2’s inhibitory effect on platelet aggregation is dependent on secondary mediators and does not interfere with collagen directly. CxD7L2 showed a low binding affinity for U-46619, the stable analog of TXA_2_ (934.58 nM), and its precursor, arachidonic acid (1083.42 nM) which might explain the high doses needed to neutralize the aggregation induced by arachidonic acid. CxD7L2 also binds serotonin and epinephrine which act as weak platelet agonists alone, but are important as they reduce the threshold concentrations of other agonists for platelet aggregation, as previously observed for the biogenic amine-binding protein from the triatomine *Rhodnius prolixus*^36^.

In contrast, we have demonstrated the novel mechanism by which CxD7L1 inhibits platelet aggregation, never reported before in the D7 protein family. CxD7L1 inhibited aggregation induced by low doses of ADP or collagen in a dose-dependent manner. Platelet aggregation induced by low doses of collagen is known to be highly dependent on ADP release from platelet granules, as platelets treated with apyrase or ADP receptor antagonists poorly respond to these agonists^37, 38^. CxD7L1 showed an inhibitory effect on aggregation triggered by the TXA_2_ pathway, as it attenuated aggregation induced by both U-46619 and arachidonic acid, the TXA_2_ precursor, which suggests that CxD7L1 interacts with TXA_2_. However, we showed CxD7L1 does not bind TXA_2_ through ITC and aggregation studies, ruling out the direct interaction between CxD7L1 and TXA_2_. It is known that aggregation through TXA_2_ is linked to ADP signaling^39^. This observation agrees with a previous description of a *R. prolixus* aggregation inhibitor 1 (RPAI-1) which binds ADP and interferes with TXA_2_ pathways^28^. Taken all together, we demonstrated that CxD7L1 inhibits platelet aggregation by sequestering ADP, which is released from platelet dense granules upon platelet activation promoting a stable platelet response^32, 33, 40^. By removing secreted ADP from the vicinity of the platelet, CxD7L1 prevents ADP from performing its role of platelet propagation.

Adenine nucleotides and derivatives play an important role in vascular biology and immunology at the mosquito bite site. ATP and ADP induce constriction of blood vessels and ADP acts as a potent mediator of platelet aggregation in mammals. Metabolism of ATP and ADP would lead to the production of AMP by apyrases that would be further metabolized to adenosine by 5-nucleotidase. Apyrases have been found in the saliva of most blood feeding arthropods studied so far^12^. The ability of CxD7L1 to scavenge ATP and ADP may compensate for the low salivary apyrase activity detected in *C. quinquefasciatus* compared to *Ae. aegypti*^41^. CxD7L1 also binds and scavenges adenosine. Although adenosine causes vasodilation and inhibits platelet aggregation, it also stimulates pain receptors and triggers pain and itch responses by inducing mast cell degranulation. Pain and itch may alert the host to the presence of a biting mosquito, preventing a successful blood meal^42^.

Arthropods underwent multiple independent evolutionary events to adapt to consume blood meals from different or new hosts. This independent evolutionary scenario has led to a great variety of salivary protein families that have acquired different functions related to blood-feeding. Gene duplication is an important mechanism for the evolution of salivary proteins. Duplication of D7 genes may have been advantageous in providing greater amounts of D7 proteins at the bite site to counteract high concentrations of host mediators^43^. Gene duplication combined with the pressure of the host hemostatic and immune responses may have led to functional divergence as observed in the D7 short proteins from *An. gambiae* and their specialization towards different biogenic amines^10^. The D7 protein family is polygenic in all Nematocera so far studied^44^. In *C. quinquefasciatus,* D7 genes are also a result of gene duplication events, given the number of genes that encode D7 proteins and their location in the genome on chromosome 3^45^. *Culex quinquefasciatus* mosquitoes are traditionally considered bird-feeders that later adapted to mammalian blood-feeding. They are increasingly recognized as important bridge vectors, vectors that acquire a pathogen from an infected wild animal and subsequently transmit the agent to a human, based on studies that examine host preference, vector/host abundance, viral infection rates, and vector competence^46^. *Culex quinquefasciatus* contain potent salivary proteins that counteract bird thrombocytes aggregation mediators such as serotonin and platelet activation factor (PAF). We have demonstrated that CxD7L2 tightly binds serotonin while Ribeiro *et al*. demonstrated that PAF phosphorylcholine-hydrolase inhibits PAF enzymatically^46^. Thrombocytes are not responsive to ADP^47, 48^, but ADP is an important mediator of platelet aggregation in mammals. We hypothesize that the novel function of ADP-binding by CxD7L1 protein has arisen from the selective pressure of mammalian hemostatic responses. This acquired D7-ADP-binding function may have provided an advantageous trait in *C. quinquefasciatus* mosquitoes that helped them to adapt to blood-feeding on mammals. *Culex tarsalis* mosquitoes prefer to feed on birds but will readily feed on mammals in the absence of their preferred host^49^. An alignment between CxD7L1 and *C. tarsalis* D7 long proteins showed that most of the residues involved in ADP binding are conserved in *C. tarsalis*, suggesting that D7 proteins that bind ADP may be widespread in the genera *Culex*. More studies are necessary to confirm this hypothesis.

In conclusion, we determined the binding capabilities of the CxD7L1 and CxD7L2 proteins and demonstrated their role in inhibiting human platelet aggregation through different mechanisms of action. We identified a novel function of ADP-binding in the well-characterized D7 protein family. Moreover, the structure of the complex CxD7L1-ADP was solved, showing a different binding mechanism for a D7 with the binding pocket located between the N-terminal and C-terminal domains whereas most D7s bind ligands within one of these two respective domains. These proteins help blood feeding in mosquitoes by scavenging host molecules at the bite site that promote vasoconstriction, platelet aggregation, itch, and pain. Accumulation of these proteins in the salivary glands of females confers an evolutionary advantage for mosquito blood feeding on mammals.

## 4. Methods

### 4.1 **Ethics statement**

Public Health Service Animal Welfare Assurance #A4149-01 guidelines were followed according to the National Institute of Allergy and Infectious Diseases (NIAID), National Institutes of Health (NIH) Animal Office of Animal Care and Use (OACU). These studies were carried out according to the NIAID-NIH animal study protocol (ASP) approved by the NIH Office of Animal Care and Use Committee (OACUC), with approval ID ASP-LMVR3.

### 4.2 Mosquito rearing and salivary gland dissection

*Culex quinquefasciatus* mosquitoes were reared in standard insectary conditions at the Laboratory of Malaria and Vector Research, NIAID, NIH (27 °C, 80% humidity, with a 12-h light/dark cycle) under the expert supervision of Andre Laughinghouse, Kevin Lee, and Yonas Gebremicale. The mosquito colony was initiated from egg rafts collected in Hilo, Hawaii, US, and maintained at NIH since 2015. Salivary glands from sugar-fed 4 to 7-day old adult mosquitoes were dissected in PBS pH 7.4 using a stereomicroscope. Salivary gland extract (SGE) was obtained by disrupting the gland wall by sonication (Branson Sonifier 450). Tubes were centrifuged at 12,000 × g for 5 min and supernatants were kept at -80 °C until use.

### 4.3 CxD7L1 and CxD7L2 gene expression pattern

*Culex quinquefasciatus* larvae (stages L1 to L4 categorized by age and size), pupae, and adults (male and female) were collected and kept in Trizol reagent (Life Technologies). Additionally, female adults were dissected, head and thorax were separated from abdomens, and independently analyzed. In all cases each sample consisted of 10 specimens. Total RNA was isolated with Trizol reagent following the manufacturer instructions (Life Technologies). cDNA was obtained with the QuantiTect Reverse Transcriptase Kit (Qiagen), from 1 µg of starting RNA. Nanodrop ND-1000 spectrophotometer was used to determine all concentrations and OD_260/280_ ratios of nucleic acids. qPCR was carried out as previously described^50^. Specific primers to target CxD7L1 and CxD7L2 genes were designed (CxD7L1-F: 5’-ACGGAAGCATGGTTTTTCAG-3’, CxD7L1-R: 5’-GGATTGCAGATTCGTCCATT-3’, CxD7L2-F: 5’-CCACGAACAACAACCATCTG-3’, CxD7L2-R: 5’-CACGCTTGATTTCATCAGGA-3’). Briefly, in a final volume of 20 µl, reaction mix was prepared with 2X SsoAdvanced Universal SYBR Green Supermix (Bio-Rad), 300 nM of each primer, and 100 ng of cDNA template. Two biological replicates were tested. All samples were analyzed in technical duplicates and non-template controls were included in all qPCR experiments as negative controls. qPCR data were manually examined and analyzed by the ΔΔCt method. ΔCt values were obtained by normalizing the data against *C. quinquefasciatus* 40S ribosomal protein S7 transcript (AF272670; CxS7-F: 5’-GTGATCAAGTCCGGCGGTGC-3’ and CxS7-R: 5’-GCTTCAGGTCCGAGTTCATCTC-3’) as the reference gene. Male adult samples were chosen as controls for the ΔΔCt values. Relative abundance of genes of interest was calculated as 2^−ΔΔCt^.

### 4.4 Cloning, expression and purification of recombinant proteins

CxD7L1 and CxD7L2 coding DNA sequences (AF420269 and AF420270) were codon-optimized for mammalian expression and synthesized by BioBasic Inc. VR2001-TOPO vectors containing CxD7L1 and CxD7L2 sequences (Vical Incorporated) and a 6x-histidine tag were transformed in One Shot TOP10 chemically competent *E. coli* (Invitrogen). FreeStyle 293-F mammalian cells were transfected with sterile plasmid DNA, prepared with EndoFree plasmid MEGA prep kit (Qiagen, Valencia, CA), at the SAIC Advance Research Facility (Frederick, MD), and supernatants were collected 72 h after transfection. Recombinant proteins were purified by affinity chromatography followed by size-exclusion chromatography, using Nickel-charged HiTrap Chelating HP and Superdex 200 10/300 GL columns, respectively.

To determine the crystal structure, recombinant CxD7L1 was produced in *E. coli*. The CxD7L1 coding DNA sequence was amplified by PCR from cDNA of *C. quinquefasciatus* salivary glands and was cloned in pET-17b plasmid and expressed in BL21 pLysS cells (Invitrogen). Protein expression was carried out as previously described^51^. Inclusion bodies were refolded using 200 mM arginine, 50 mM Tris, 1 mM reduced glutathione, 0.2 mM oxidized glutathione, 1 mM EDTA, pH 8.0. Bacterial CxD7L1 was purified by size exclusion chromatography, using a HiPrep 16/60 Sephacryl S-100 HR column, followed by cation exchange chromatography with a HiPrep SP FF 16/10 column. A last step of analytical size exclusion chromatography was performed using a Superdex 200 10/300 GL column with 25 mM Tris, 50 mM NaCl pH 7.4. All HPLC columns were obtained from GE Healthcare Life Science, Piscataway, NJ. All purified proteins were separated in a 4-20% NuPAGE Tris-glycine polyacrylamide gel and visualized by Coomassie stain. Protein identity was verified by Edman degradation at the Research Technologies Branch, NIAID, NIH.

### 4.5 Polyclonal antibody production

Polyclonal antibodies against CxD7L1 and CxD7L2 were raised in rabbits. Immunization of rabbits was carried out in Noble Life Science facility according to their standard protocol (http://www.noblelifesci.com/preclinical-drug-development/polyclonal-antibody-production/). Rabbit sera were shipped to our laboratory where purification of IgG was performed by affinity chromatography using a 5-ml HiTrap protein A HP column following manufacturer’s instructions (GE Healthcare, Piscataway, NJ). Purified IgG protein concentration was determined by Nanodrop ND-1000 spectrophotometer. Additionally, antibodies against *C. quinquefasciatus* salivary gland extract were raised in rabbits. Levels of specific antibodies were determined by ELISA according to Chagas *et al*.^52^

### 4.6 Western blot

*Culex quinquefasciatus* salivary gland extracts (2.5 µg) and 100 ng of CxD7L1 and CxD7L2 were separated by NuPAGE. Proteins were transferred to a nitrocellulose membrane (iBlot, Invitrogen) that was blocked overnight at 4 °C with blocking buffer: TBS containing 5% (w/v) powdered non-fat milk. Purified anti-CxD7L1 and anti-CxD7L2 IgG antibodies were diluted in blocking buffer (0.5 µg/ml) and incubated for 90 min. Goat anti-rabbit conjugated to alkaline phosphatase (Sigma) diluted in blocking buffer (1:10,000) was used as a secondary antibody and immunogenic bands were developed by the addition of BCIP/NBT substrate (Promega). The reaction was stopped with distilled water.

### 4.7 Immunolocalization of CxD7L1 and CxD7L2

*Culex quinquefasciatus* salivary glands were dissected in PBS, transferred to a welled plate, and fixed with 4% paraformaldehyde (Sigma) for 30 min at room temperature. Tissues were washed 3 times for 10 min each with 1x PBS to remove paraformaldehyde and then blocked with 2% BSA, 0.5% Triton X-100, 1x PBS pH 7.4 overnight at 4 °C. Glands were washed 3 times with PBS to remove Triton X-100 and were transferred to clean wells to which 200 μl of 1 μg/ml pre-adsorbed antibodies against either CxD7L1 or CxD7L2 (raised in rabbits and diluted 1:1000 in 2% BSA 1x PBS) were added. Glands incubated in 2% BSA 1x PBS served as a negative control. Plate wells were covered and incubated overnight at 4 °C. Primary antibodies were removed by 3 washes with 2% BSA 1x PBS and incubated with 2 μg/ml anti-rabbit IgG Alexa Fluor 594 (Thermo Fisher) for 2 h in the dark at 4°C. Conjugate was removed by 3 additional washes with 1x PBS. DNA was stained with 1 µg/mL DAPI (Sigma D9542) and actin with 0.04 µg/mL Phalloidin Alexa 488 (Invitrogen) for 20 min. Glands were washed three times with PBS and transferred to glass slides containing droplets of PBS. PBS was removed without drying the glands, and tissues were mounted using a coverslip coated with 25 µl Prolong Gold mounting medium. Slides were covered and left to dry at room temperature and then stored at 4 °C. Bright field and fluorescent images were acquired in a Leica Confocal SP8 microscope with a 63x objective using Navigator tool. Images were processed with Imaris software version 9.2.1 and postprocessing was carried out in Fiji ImageJ for representative purposes.

### 4.8 Isothermal titration calorimetry (ITC)

Thermodynamic binding parameters of CxD7L1 and CxD7L2 to several pro-hemostatic ligands were tested using a Microcal VP-ITC microcalorimeter. The panel of substances tested included several nucleosides/nucleotides or derivates (ATP, ADP, 5’-AMP, 3’-AMP, cyclic AMP, adenosine, GTP, TTP, inosine, sodium polyphosphate, Sigma-Aldrich), biogenic amines (epinephrine, norepinephrine, histamine, serotonin, Sigma-Aldrich), and pro-inflammatory/pro-hemostatic lipid compounds (LTB_4_, LTC_4_, LTD_4_, LTE_4_, arachidonic acid, and the stable analog of TXA_2_: U-46619, Cayman Chemicals). Ligands and protein solutions were prepared in 20 mM Tris-HCl pH 7.4, 150 mM NaCl (TBS) at 30 and 3 µM, respectively. Lipids ligands were prepared by evaporating the ethanol or chloroform solvent to dryness under a stream of nitrogen. Lipid ligands were further dissolved in TBS and sonicated for 10 min (Branson 1510) to ensure dissolution. Lipid ligands were used at 50 µM of ligand and 5 µM of protein. Injections of 10 µl of ligand were added to the protein samples contained in the calorimeter cell at 300 sec intervals. Experiments were run at 30 °C. Thermodynamic parameters were obtained by fitting the data to a single-site binding model in the Microcal Origin software package. For saturation studies, CxD7L2 protein was pre-incubated with 50 µM serotonin for 30 min and titrated with LTD_4_.

### 4.9 CxD7L1 Crystallization, data collection and structure determination

Purified protein was incubated overnight at 4°C with 1.2 times molar excess of ADP. Crystals were obtained using the hanging drop-vapor diffusion method with 0.01 M Zinc sulfate heptahydrate, 0.1 M MES monohydrate pH 6.5, and 25% v/v Polyethylene glycol monomethyl ether 550 (Crystal Screen 2, Condition 27, Hampton Research).

For data collection the crystals were rapidly soaked in the mother liquor solution (the crystallization buffer described above) supplemented with 25% glycerol and flash frozen in a nitrogen gas stream at 95 K. Data were collected at beamline 22BM at the Advanced Photon Source, Argonne National Laboratory equipped with 10Hz Rayonix MX300HS detector. A crystal that diffracted to 1.97 Å resolution with cell dimensions (in Å) of a =76.66, b =84.32, and c =132.07 and belonged to the orthorhombic space group I212121 (Table 2) was used to collect a data set. The data were processed, reduced and scaled with XDS^53^. The structure of CxD7L1 was determined by molecular replacement using Phaser^54^ by employing separate, manually constructed search models for the N-terminal and C-domains based on the crystal structure of *Anopheles stephensi* AnStD7L1 (PDB ID: 3NHT). The final model of CxD7L1 was constructed by iterative manual tracing of the chain using the program Coot^55^ after each cycle of refinement with stepwise increase in the resolution using Phenix^56^. All structural figures were produced with PyMOL (PyMOL molecular graphics system, version 1.7.4; Schrödinger, LLC) and UCSF Chimera (Resource for Biocomputing, Visualization, and Informatics at the University of California, San Francisco, with support from NIH P41-GM103311)^57^.

### 4.10 Platelet aggregation assay

Platelet rich plasma (PRP) was obtained from normal healthy donors on the NCI IRB approved NIH protocol 99-CC-0168, “Collection and Distribution of Blood Components from Healthy Donors for In Vitro Research Use.” Research blood donors provide written informed consent, and platelets were de-identified prior to distribution. Platelet aggregation was measured using an aggregometer (Chrono-Log Corporation). Briefly, 300 μL of PRP, diluted 1:3 to approximately 250,000 platelets/uL in Hepes-Tyrode’s buffer (137 mM NaCl, 27 mM KCl, 12 mM NaHCO_3_, 0.34 mM sodium phosphate monobasic, 1 mM MgCl_2_, 2.9 mM KCl, 5 mM Hepes, 5 mM glucose, 1% BSA, 0.03 mM EDTA, pH 7.4) were pre-stirred in the aggregometer for 1 min to monitor pre-aggregation effects. Different concentrations of recombinant proteins or TBS as negative control were added to the PRP before adding the agonists. Aggregation agonists used in our studies included native collagen type I fibrils from equine tendons, convulxin, ADP, U-46619, arachidonic acid, serotonin, epinephrine, or combination of agonists. Their concentrations are specified in the figure captions. Technical duplicates were performed.

## Acknowledgements

We thank Kevin Lee, Andre Laughinghouse, and Yonas Gebremicale for excellent mosquito rearing and Van My Pham for salivary glands dissection. We also thank John Andersen and Jose Ribeiro for relevant scientific discussion and Thrity Avary from Chrono-Log Corporation for technical assistance with platelet aggregation studies. The authors thank Bradley Otterson, NIH Library Writing Center, for manuscript editing assistance. This research was supported by the Intramural Research Program of the NIH/NIAID (AI001246-01).

---

**Supplementary Fig. 1.**
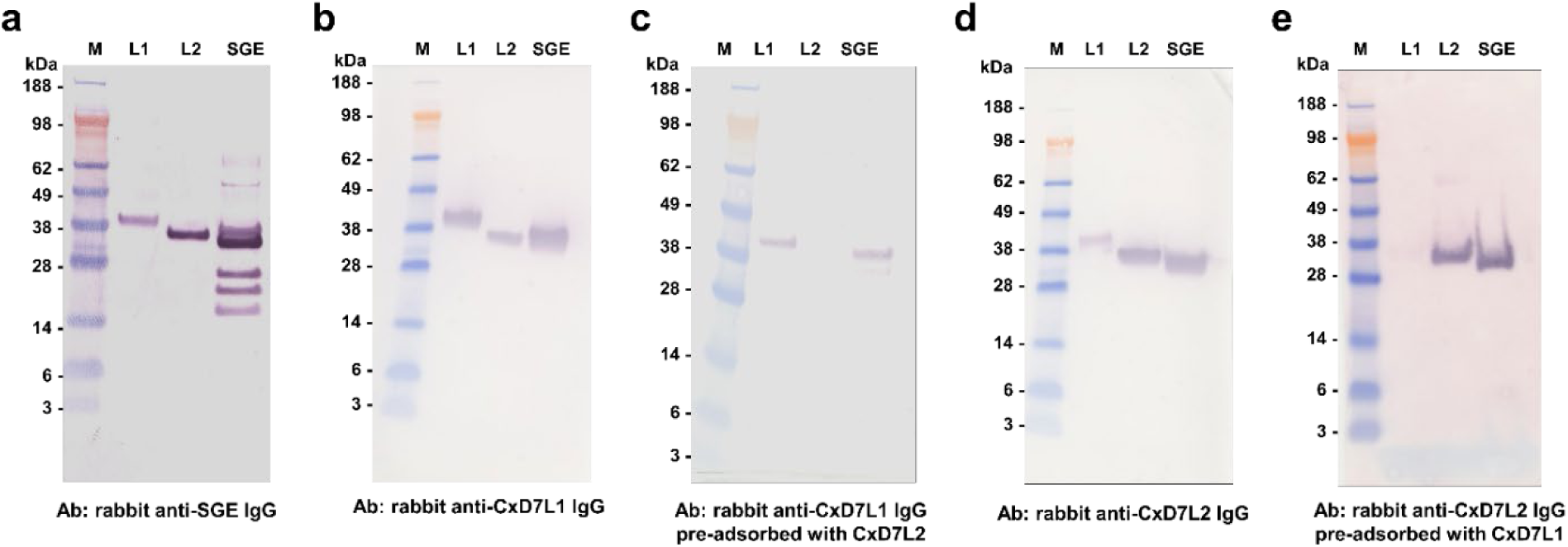
Recognition of recombinant CxD7L1 and CxD7L2 by IgG antibodies raised in rabbits. (a) Purified IgG from serum of a rabbit immunized with salivary gland extract (SGE) from *Culex quinquefasciatus* recognized the recombinant proteins CxD7L1 and CXD7L2 (100 ng) and other protein bands from the salivary gland extract (2.5 µg). **(b)** Purified IgG from serum of a rabbit immunized with CxD7L1 protein recognized CxD7L1 recombinant protein (100 ng) and a band of similar molecular weight in the SGE (2.5 µg). It also cross-reacted with CxD7L2. **(c)** Purified IgG from serum of a rabbit immunized with CxD7L1 protein and pre-adsorbed with CxD7L2 specifically recognized CxD7L1 recombinant protein (100 ng) and a band of similar molecular weight in the SGE (2.5 µg). **(d)** Purified IgG from serum of a rabbit immunized with CxD7L2 protein recognized CxD7L2 recombinant protein (100 ng) and a band of similar molecular weight in SGE (2.5 µg). It also cross-reacted with CxD7L1 **(e)** Purified IgG from serum of a rabbit immunized with CxD7L2 protein and pre-adsorbed with CxD7L1 specifically recognized CxD7L2 recombinant protein (100 ng) and a band of similar molecular weight in the SGE (2.5 µg). No cross-reactivity between anti-CxD7L1 IgG and anti-CxD7L2 IgG was observed after anti-CxD7L1 was pre-adsorbed with CxD7L2 and anti-CxD7L2 was pre-adsorbed with CxD7L1. Anti-Culex SGE IgG antibodies were used at 1 µg/ml and IgG antibodies against recombinant proteins were used at 0.5 µg/ml. Goat anti-rabbit IgG AP (1:10,000 dilution, Sigma) was used as a secondary antibody. SeeBlue Plus2 Pre-stained was used as the protein standard (M).

**Supplementary Fig. 2.**
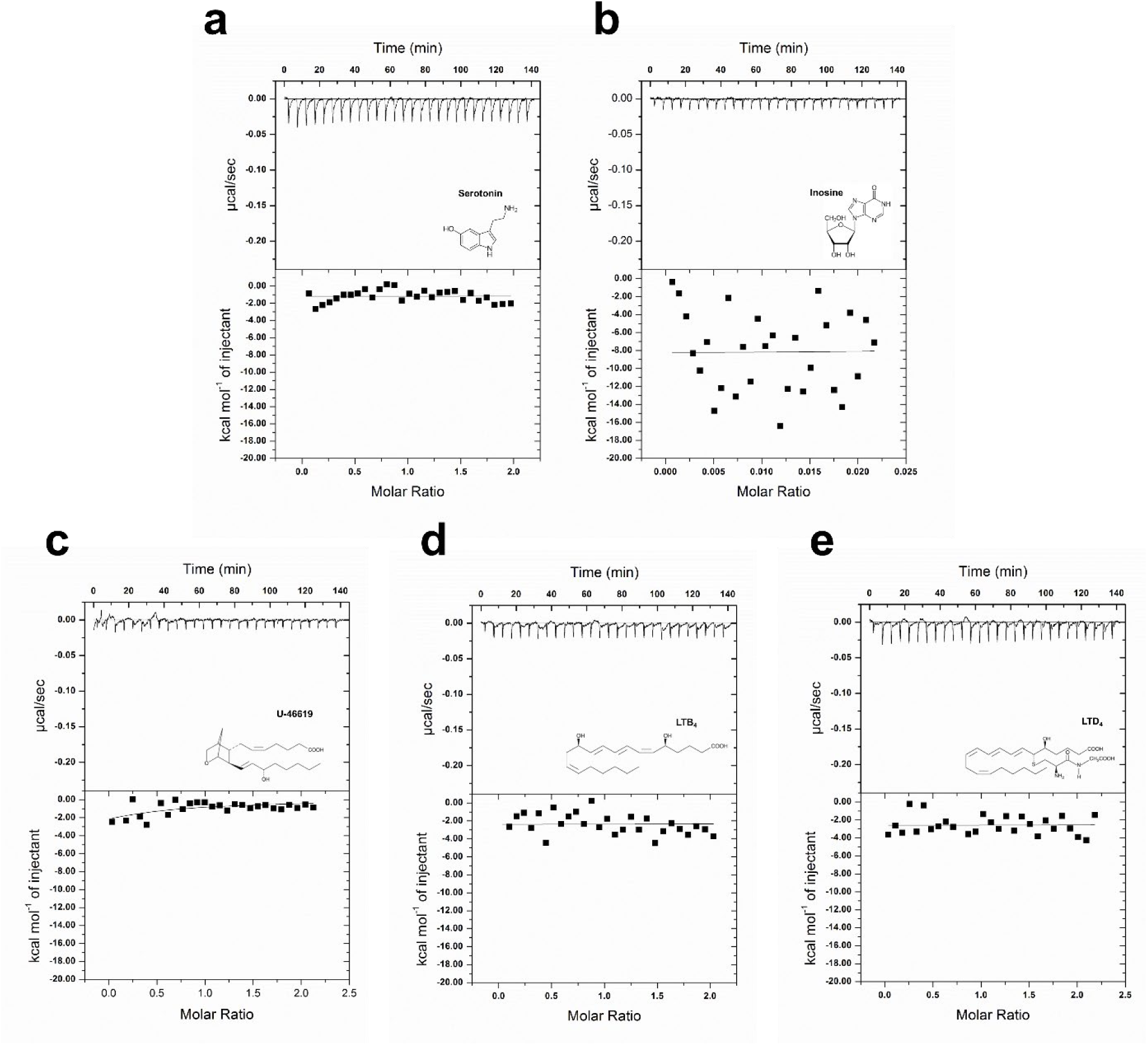
Isothermal titration calorimetry studies of CxD7L1. Binding experiments were performed on a VP-ITC microcalorimeter. **(a)** 30 µM serotonin or **(b)** 30 µM inosine were titrated with 3 µM of CxD7L1. For TXA_2_ analog U-46619 **(c)** and leukotrienes LTB_4_ **(d)** and LTD_4_ **(e)** protein and ligand were prepared at 5 µM and 50 µM, respectively. Assays were performed at 30 °C. The upper curve in each panel shows the measured heat for each injection, while the lower graph shows the enthalpies for each injection and the fit to a single-site binding model for calculations of thermodynamic parameters. The insets show the names and chemical formulas for these compounds.

**Supplementary Fig. 3.**
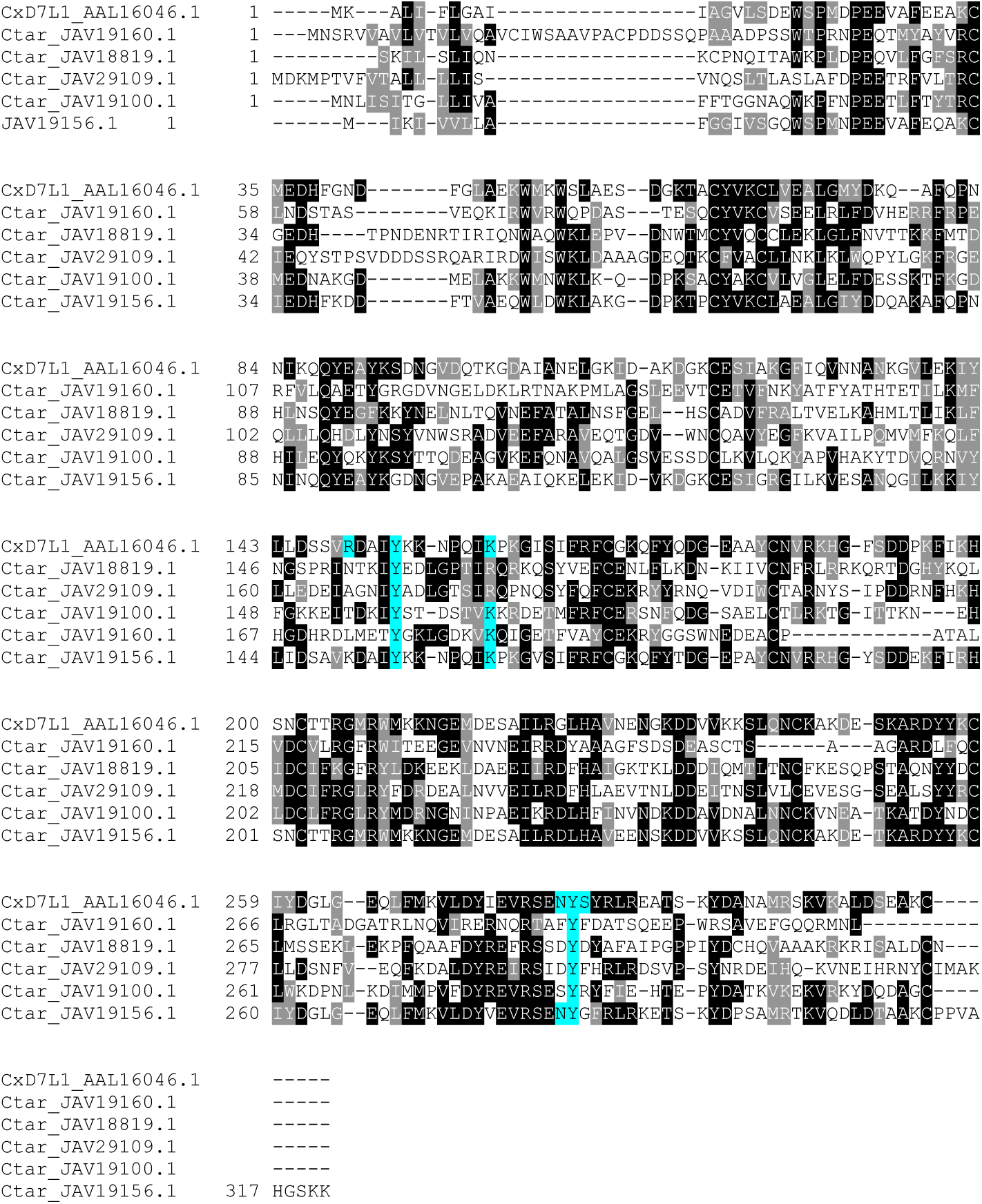
Multiple sequence alignment of *Culex quinquefasciatus* CxD7L1 and *Culex tarsalis* D7 long proteins. Eighteen D7 homologs from *C. tarsalis* were retrieved from NCBI database after a tBLAST search using the CxD7L1 protein as the query sequence. The database used was the Transcriptome Shotgun Assembly, BioProject PRJNA360148. *Culex tarsalis s*equences with E value lower than 4e-10 were chosen (N = 10) and clustered by cd-hit software^25^ where sequence identity cut-off was set at 0.85. CxD7L1 (AAL16046) and 5 representative of *C. tarsalis* D7 long protein homologs (JAV19160, JAV18819, JAV29109, JAV19100, and JAV19156) were aligned with Clustal Omega and refined using BoxShade server. Black background shading represents identical amino acids (50% of sequences must agree for shading) while grey shading designates similar amino acids. Residues highlighted in blue indicate conserved amino acids involved in ADP binding.

